# Prenatal alcohol and cannabinoid co-exposure increases ethanol-seeking in adult offspring and produces sex-dependent medial prefrontal cannabinoid receptor 1 expression

**DOI:** 10.64898/2026.09.21.753297

**Authors:** Siara K. Rouzer, Abigail Bowring, Aisley George, McKay Domen, Emma Labbe, Marisol Palacios, Rajesh C. Miranda

## Abstract

**Background:** Alcohol and cannabinoids are frequently used during pregnancy, however their combined effects on offspring outcomes are currently under-investigated. In this study, co-exposure was compared to single-drug and drug-free exposures for influences on adult alcohol-seeking and cannabinoid system proteomics.

**Methods:** Pregnant C57BL/6J mice received ethanol vapor (ALC), CP-55,940 (0.75 mg/kg; CB), both (ALC+CB), or neither on gestational days 12–15. Beginning on Postnatal Day 200, male and female offspring underwent home-cage ethanol drinking (N=108) and operant self-administration, progressive-ratio, extinction, and reinstatement testing (N=96). CNR1 and CNRIP1 protein abundance was quantified in the medial prefrontal cortex (mPFC) and dorsomedial striatum (N=57) and compared to within-subject drinking behaviors.

**Results:** Prenatal CB exposure increased cumulative home-cage ethanol intake, primarily in males. At 40% ethanol under progressive-ratio schedules, ALC+CB males consumed more than all other male groups, whereas all exposed female groups consumed more than controls. ALC+CB males also showed greater extinction responding than all other male groups and greater reinstatement intake relative to baseline than CON and ALC males. ALC+CB increased mPFC CNR1 in males relative to all other groups, whereas CB and ALC+CB reduced mPFC CNR1 in females. Higher mPFC CNR1 was associated with fixed-ratio and 40% progressive-ratio intake, extinction, and reinstatement exclusively in males.

**Conclusions:** Prenatal co-exposure produced a persistent male phenotype of enhanced ethanol seeking that was distinct from either exposure alone and accompanied by elevated mPFC CNR1. This behavioral and molecular convergence justifies future investigations manipulating corticostriatal cannabinoid signaling to attenuate exposure-induced ethanol consumption in late adulthood.

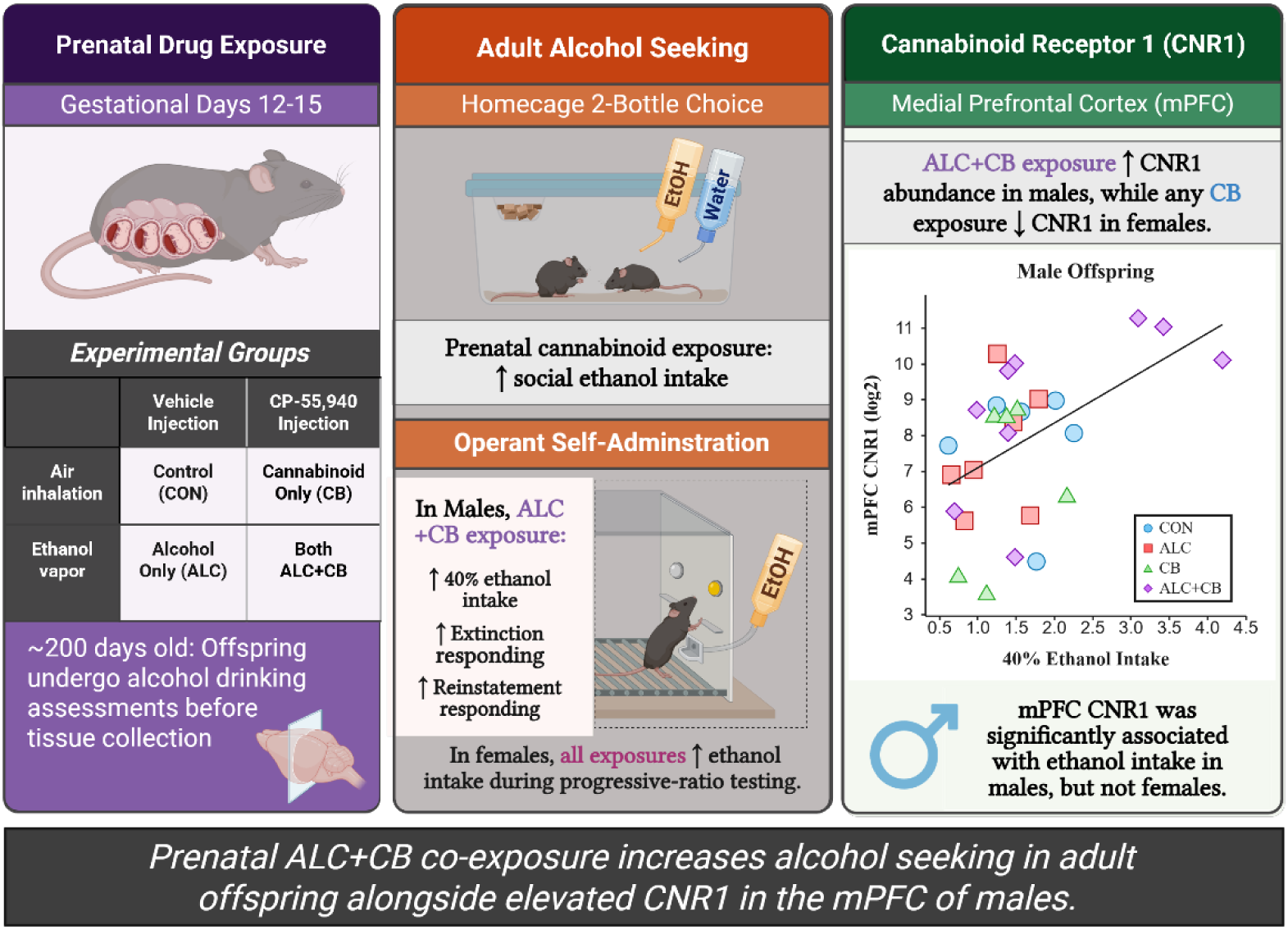

## Introduction

Alcohol use disorder (AUD) is one of the most prevalent mental health disorders worldwide, affecting an estimated 4–5% of the population, including ∼8% of men and ∼1.5% of women [1, 2]. During reproductive years, the prevalence of AUD increases the likelihood that offspring will be exposed to alcohol before birth. In the United States, approximately 10% of pregnant individuals report alcohol use in the past 30 days, with binge drinking reported by 4.5% [3]. Although AUD is moderately heritable [4] and genome-wide association studies have identified multiple associated genetic loci [5], genetic susceptibility does not fully account for its familial transmission.

Prenatal alcohol exposure (PAE) represents a distinct developmental pathway through which alcohol use during pregnancy may increase offspring vulnerability to later alcohol misuse, potentially contributing to its transmission across generations. A substantial body of preclinical and clinical research suggests that PAE increases vulnerability to later-life alcohol misuse. Clinical studies associate PAE with alcohol-related problems, alcohol disorders, and multiple AUD symptoms by early adulthood [6–8]. Preclinical studies similarly demonstrate increased ethanol consumption following PAE, although the timing and magnitude of these effects vary by sex, age, and ethanol concentration [9–12].

Since AUD commonly co-occurs with other substance use disorders [13], the consequences of PAE may be modified by prenatal exposure to other psychoactive substances. Alcohol and cannabis are frequently co-used by individuals of reproductive age [14, 15], creating the potential for combined prenatal exposure during unplanned pregnancies [16]. Co-use is associated with sexual risk-taking behaviors [17, 18], higher incidence of substance use disorders [19], and increased risk of alcohol misuse [20]. Clinical and preclinical studies indicate bidirectional pharmacokinetic interactions during alcohol-cannabinoid co-exposure, with alcohol increasing circulating THC and 11-OH-THC concentrations and cannabinoids increasing blood alcohol concentrations [14, 21–23]. These bidirectional pharmacokinetic interactions are consistent with increased systemic exposure during co-use.

Despite the widespread co-use of these substances, few studies have examined alcohol-directed behavior following combined prenatal alcohol and cannabinoid exposure. Ornelas et al. found that prenatal exposure to the synthetic cannabinoid CP-55,940, alone or with alcohol, increased adolescent ethanol intake in both sexes and adult voluntary ethanol consumption in females, with combined exposure also increasing open-field activity in males [24]. Compared to single-drug exposure, prenatal alcohol and cannabinoid co-exposure has also reduced fetal growth and viability [25, 26], impaired motor coordination and increased locomotor activity [26, 27], and reduced fetal cerebral blood flow [28]. However, it remains unknown whether prenatal co-exposure alters ethanol seeking in middle adulthood, whether these effects depend on social context, and which persistent neurobiological changes are associated with these behaviors.

The endocannabinoid system is a strong candidate mechanism linking prenatal co-exposure to altered ethanol-directed behavior. This system regulates neurodevelopment and ethanol reinforcement and is persistently altered by developmental exposure to alcohol and cannabinoids [14, 29, 30]. Within corticostriatal circuitry, cannabinoid receptor 1 (CNR1) is expressed on presynaptic terminals, where its activation suppresses glutamate release [31], and contributes to ethanol reinforcement and seeking [29]. Voluntary ethanol intake also strengthens glutamatergic transmission from the medial prefrontal cortex (mPFC) onto direct-pathway neurons in the dorsomedial striatum (DMS), whereas bidirectional manipulation of this plasticity correspondingly increases or decreases ethanol seeking [32]. CNR1 signaling is modulated by cannabinoid receptor interacting protein 1 (CNRIP1), which regulates receptor signaling and downregulation [33]. Together, corticostriatal CNR1/CNRIP1 demonstrate promise as mechanisms associating prenatal alcohol and cannabinoid co-exposure with persistent ethanol-directed behavior.

Our current study builds on this literature by examining how prenatal co-exposure to alcohol and CP-55,940 affects ethanol drinking and seeking in adult mouse offspring.

Specifically, we assessed ethanol-directed behavior in social and reinforced contexts in mature adulthood, quantified CNR1 and CNRIP1 protein expression in the mPFC and DMS, and evaluated associations between protein expression and behavior. Given prior sex- and exposure-dependent outcomes, we hypothesized that co-exposure would produce enduring, sex-specific increases in ethanol drinking and seeking. We further tested whether CNR1 and CNRIP1 expression differed by sex and exposure in association with ethanol-seeking behaviors.

## Methods & Materials

### Experimental animals and study design

Adult male and female C57BL/6J offspring were selected from our previously characterized prenatal exposure cohort [26], where pregnant dams were randomly assigned to control (CON), alcohol (ALC), cannabinoid (CB), or combined alcohol and cannabinoid (ALC+CB) conditions. From gestational days 12–15, dams received the synthetic CB1/CB2 receptor agonist CP-55,940 (0.75 mg/kg, i.p.) or vehicle immediately before a 30-min exposure to ethanol vapor or room air. The present cohort represented 22 litters. All procedures were approved by the Texas A&M University Institutional Animal Care and Use Committee (Protocol No. 2021-0331).

Offspring completed home-cage ethanol drinking, operant acquisition and ethanol fixed-ratio self-administration, progressive-ratio testing, extinction, reinstatement, and tissue collection in a fixed sequence during middle adulthood (Fig. 1). Home-cage drinking was assessed in 108 offspring housed across 46 same-sex, exposure-matched cages, with cage treated as the experimental unit. Operant assessments included 96 offspring, and protein measurements were obtained from 57 offspring. Expanded descriptions of cohort composition, experimental apparatus and procedures, outcome calculations, quality-control criteria, and statistical models are provided in the corresponding sections of the Supplemental Methods.

**Figure 1.**
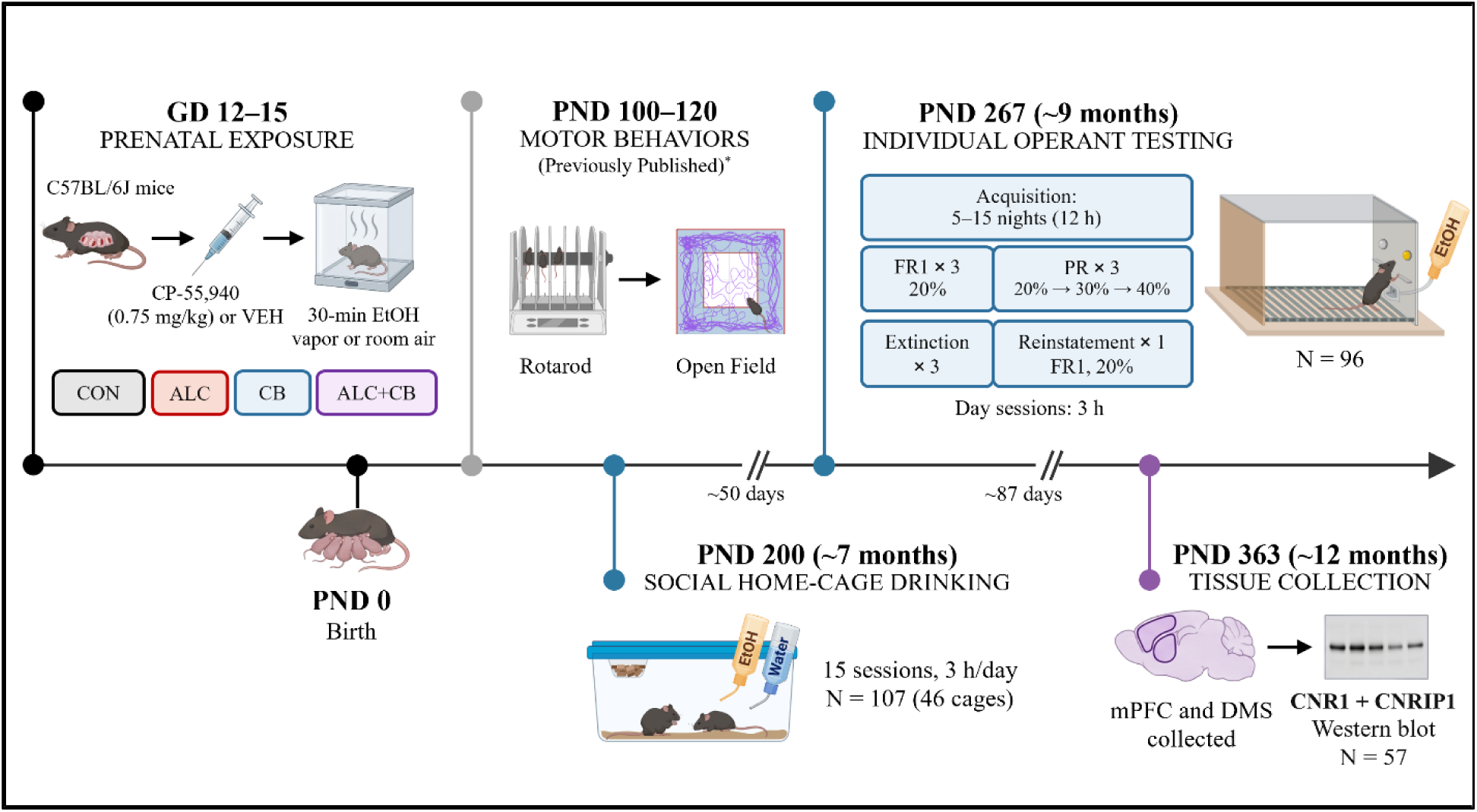
Experimental timeline and study design. Pregnant C57BL/6J mice received CP-55,940 (0.75 mg/kg) or vehicle (VEH) and 30 min of ethanol (EtOH) vapor or room air daily from gestational day (GD) 12–15, producing four prenatal exposure groups: control (CON), alcohol (ALC), cannabinoid (CB), and combined alcohol and cannabinoid exposure (ALC+CB). Birth was designated postnatal day (PND) 0. Offspring underwent rotarod and open-field testing at PND 100–120. These motor-behavior assessments were reported previously (Rouzer et al., 2025; doi:10.1016/j.dadr.2025.100356). Beginning at approximately PND 200, offspring completed 15 social home-cage drinking sessions conducted for 3 h/day (N=107 offspring in 46 same-sex cages). Approximately 50 days later, offspring began individual operant ethanol testing at PND 267. Operant testing included 5–15 overnight acquisition sessions (12 h/session), three fixed-ratio 1 (FR1) sessions with 20% EtOH, three progressive-ratio (PR) sessions with sequential access to 20%, 30%, and 40% EtOH, three extinction sessions, and one FR1 reinstatement session with 20% EtOH. Daytime operant sessions lasted 3 h (N=96). Approximately 87 days after operant testing, medial prefrontal cortex (mPFC) and dorsomedial striatum (DMS) tissue was collected at PND 363 for Western blot quantification of CNR1 and CNRIP1 protein (N=57). Approximate ages and intervals are shown.

### Voluntary home-cage ethanol drinking

Voluntary ethanol drinking was assessed across 15 weekday sessions conducted over 19 calendar days. Each session lasted 3 h and occurred during the light phase. During sessions 1–5, mice received one tube containing 20% ethanol. During sessions 6–15, a second tube containing water was added to assess ethanol-to-water preference. Ethanol intake was calculated from the change in ethanol-tube weight, ethanol concentration, and average body weight of the cage occupants. Cumulative intake was calculated across all 15 sessions. Preference was defined as the natural logarithm of the ratio of cumulative ethanol-solution volume to cumulative water volume across sessions 6–15, with 0 representing equal consumption. The cage served as the experimental unit.

### Operant ethanol self-administration, extinction, and reinstatement

Operant testing used a lever-based modification of the ‘CUP’ paradigm [34]. The right lever was active for all animals, whereas the left lever was inactive. During reinforced sessions, each active-lever press delivered 18 µL of ethanol, illuminated the stimulus light above the active lever, and initiated a 5-s timeout. Acquisition consisted of 12-h overnight fixed-ratio 1 (FR1) sessions with 20% ethanol. Acquisition required at least 100 total lever presses and at least 70% active-lever responding during each of three consecutive sessions. Training continued until both criteria were met or for a maximum of 15 sessions.

Following acquisition, mice completed three 3-h FR1 sessions with 20% ethanol, three 3-h progressive-ratio sessions with 20%, 30%, and 40% ethanol, three 3-h extinction sessions, and one 3-h reinstatement session. Breakpoint was defined as the final progressive-ratio response requirement completed. During extinction, lever presses produced neither ethanol nor stimulus-light illumination. Reinstatement used the previous 20% ethanol FR1 contingency. Ethanol intake was calculated from the change in syringe volume after subtraction of ethanol recovered from the receptacle and was normalized to body weight.

### Blood ethanol collection

All subjects underwent tail-blood collection immediately after the final 3-h home-cage drinking, FR1 session, and reinstatement sessions. However, because ethanol detection was low and variable despite independently verified consumption (see Supplemental Methods), post-session blood ethanol concentrations were not reported as reliable consumption outcomes.

### Tissue collection and Western blot analyses

Following ketamine/xylazine anesthesia and secondary decapitation without perfusion, the bilateral medial prefrontal cortex (mPFC) and dorsomedial striatum (DMS) were dissected from fresh tissue with reference to the Allen Mouse Brain Atlas. Tissue was sonicated in RIPA buffer containing protease inhibitor and centrifuged at 20,800 × g for 10 min at 4°C. Protein concentration was determined using a bicinchoninic acid assay. For each sample, 2 µg of protein was separated on a 12% Bis-Tris gel and transferred to a PVDF membrane.

Total protein was visualized using REVERT Total Protein Stain (LI-COR Biosciences). Membranes were co-incubated with rabbit anti-CNR1 (Abcam, catalog no. ab259323; 1:100) and rabbit anti-CNRIP1 (Proteintech, catalog no. 16827-1-AP; 1:100), followed by IRDye 800CW goat anti-rabbit IgG secondary antibody (LI-COR Biosciences; 1:14,300). Fluorescent signals were acquired using an Odyssey CLx and analyzed in Image Studio version 5.2. CNR1 and CNRIP1 immunoreactivity was quantified at approximately 53 and 20 kDa, respectively. Signals were normalized sequentially within and among membranes using REVERT total-protein signals. Uncropped immunoblots and corresponding total-protein stains are provided in Figure S1.

### Statistical analyses and scientific rigor

Subjects were identified by numeric codes and tail colors that concealed prenatal exposure history. Operant chamber assignments were randomized, and experimenters remained blinded throughout behavioral testing, tissue collection, Western blotting, and protein quantification. Hand-recorded home-cage and operant syringe measurements were independently transcribed by two experimenters and electronically compared. Statistical analyses were conducted using IBM SPSS Statistics version 29.0.2.0 and GraphPad Prism version 11. Prenatal ALC and CB exposure were treated as separate factors in factorial analyses. Initial models included Sex, with sex-stratified follow-up analyses conducted for significant or trend-level effects involving Sex and to characterize established sex differences in ethanol-directed behavior. Four-level exposure analyses were used when needed to distinguish CON, ALC, CB, and ALC+CB groups. Šídák-adjusted comparisons followed significant or trend-level omnibus effects unless otherwise specified. Potential outliers were evaluated separately for each outcome using the ROUT procedure with Q = 1%.

Repeated home-cage and operant outcomes were evaluated using linear mixed-effects models with random intercepts for cage or subject, respectively, and the relevant repeated factor of day or ethanol concentration. Home-cage preference, acquisition, reinstatement, and normalized outcomes were evaluated using factorial models. Preference was compared with 0, and normalized extinction and reinstatement outcomes were compared with 100%, using corrected one-sample tests. Protein-expression values were transformed as ln(expression + 1) and analyzed with Region as a within-subject factor. Protein-behavior relationships were assessed separately by sex using multivariate general linear models followed by corrected correlations. Statistical significance was defined as *p* < .05, and .05 ≤ *p* < .10 was reported as a statistical trend. Partial eta squared (ηp²) is reported as effect size.

## Results

Comprehensive model outputs, secondary outcome analyses, and sensitivity analyses are reported in the Supplementary Results and Tables S1–S37.

### Prenatal cannabinoid exposure increased voluntary social drinking, particularly in male offspring

Across 15 home-cage drinking sessions, cumulative cage-level ethanol intake was higher in females than males [*F*(1,38.05)=7.90, *p*=.008] (Fig. 2A). Consistent with prior research in C57BL/6J mice [35, 36], control females consumed more ethanol per day than control males (1.99 versus 1.10 g/kg/day), [nested *t*(28)=6.65, *p*<.001]. Prenatal CB exposure produced a trend toward increased cumulative intake [*F*(1,38.05)=3.42, *p*=.072], and interacted with Drinking Day [*F*(14,516.10)=3.05, *p*<.001], (Fig. 2B; Table S1).

**Figure 2.**
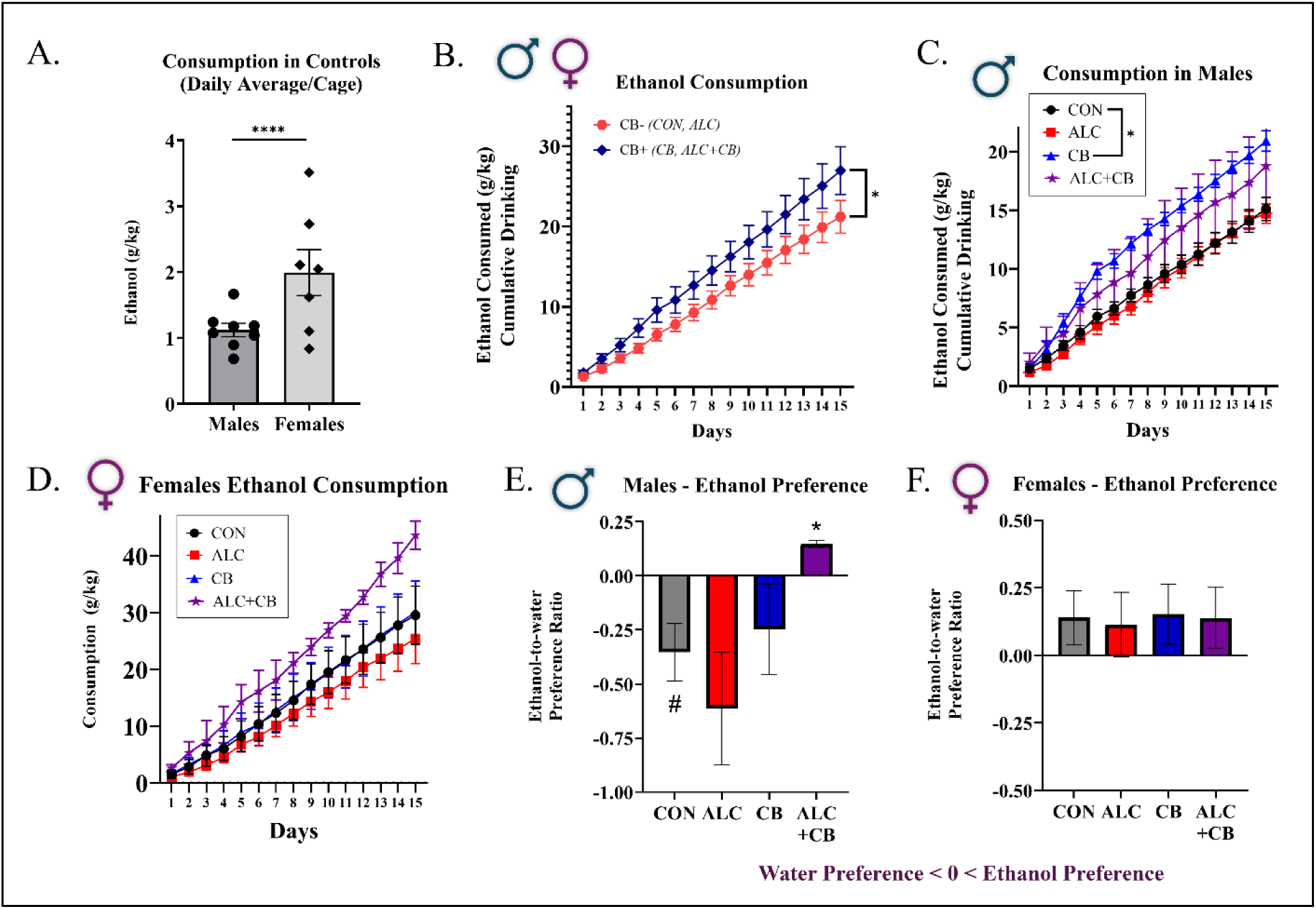
Home-cage ethanol consumption and preference following prenatal alcohol and cannabinoid exposure. (A) Mean daily ethanol consumption among control offspring, demonstrating greater consumption in females than males. Symbols represent individual cages, and bars show mean ± SEM. (B) Cumulative ethanol consumption across the 15-day assessment in offspring without prenatal cannabinoid exposure (CB−: CON and ALC) and with prenatal cannabinoid exposure (CB+: CB and ALC+CB), collapsed across sex. (C, D) Cumulative ethanol consumption across the 15-day assessment in male (C) and female (D) offspring from the CON, ALC, CB, and ALC+CB groups. (E, F) Mean natural-log-transformed ethanol-to-water consumption ratio across the 10-day preference period in male (E) and female (F) offspring. A value of 0 indicates equal ethanol and water consumption, negative values indicate greater water consumption, and positive values indicate greater ethanol consumption. Unless otherwise indicated, data are presented as mean ± SEM, with cage as the statistical unit. The control-group sex comparison in (A) was evaluated using a nested *t*-test with offspring nested within cage. Longitudinal ethanol intake was evaluated using linear mixed-effects models with Day as a repeated factor and cage as a random intercept. Preference ratios in (E) and (F) were compared with 0 using two-tailed one-sample *t*-tests with Holm correction across exposure groups within each sex. CON, control; ALC, prenatal alcohol exposure; CB, prenatal cannabinoid exposure; ALC+CB, combined prenatal alcohol and cannabinoid exposure. #*p* < .10; \**p* < .05; \*\*\*\**p* < .0001.

In males, CB-exposed offspring consumed more ethanol than non-CB-exposed offspring, [*F*(1,19.78)=9.78, *p*=.005, ηp²=.331], with greater cumulative intake from Days 4 through 15. CB-only males also consumed more than CON males from Days 7 through 15 (all Šídák-adjusted *ps* ≤ .049; Fig. 2C; Table S2). ALC exposure interacted with Day [*F*(14,249.55)=2.26, *p*=.006], although ALC-exposed and non-ALC-exposed males did not differ on any individual day. Females showed no exposure main effects; however, CB exposure produced a trend-level interaction with Day [*F*(14,265.09)=1.60, *p*=.080], and CB-exposed females had greater cumulative intake on Day 15 (*p*=.045; Fig. 2D; Table S3).

Across the preference phase, females had higher ethanol-to-water consumption ratios than males, [*F*(1,38.19)=9.89, *p*=.003, ηp²=.206]. CB exposure produced a trend toward increasing ethanol-to-water ratios independent of sex, [*F*(1,38.19)=2.95, *p*=.094], and in males [*F*(1,19.04)=4.26, *p*=.053]. Relative to equal liquid preference, ALC+CB males consumed proportionally more ethanol than water, [*t*(2)=9.38, Holm-adjusted *p*=.044], whereas no female group preferred one liquid over the other (Fig. 2E–F; Tables S4–S5).

### Prenatal exposures produced sex- and task-dependent increases in operant ethanol self-administration

Days required to reach the acquisition criterion were not affected by Sex, ALC, or CB exposure, but differed as a function of Sex × ALC exposure [*F*(1,88)=6.74, *p*=.011], although adjusted within-sex comparisons did not reach significance (Table S6). During FR1 self-administration, females consumed more ethanol than males [*F*(1,85.54)=7.04, *p*=.009, ηp²=.076], and a Sex × ALC × CB interaction was observed [*F*(1,85.54)=4.36, *p*=.040; Fig. 3A]. CB exposure increased FR1 intake in females *[F*(1,41.45)=4.09, *p*=.050, ηp²=.090], with CB-only females consuming more than CON females (*p* = .015). No exposure effects on FR1 intake were detected in males (Fig. 3B–C; Tables S7–S9).

**Figure 3.**
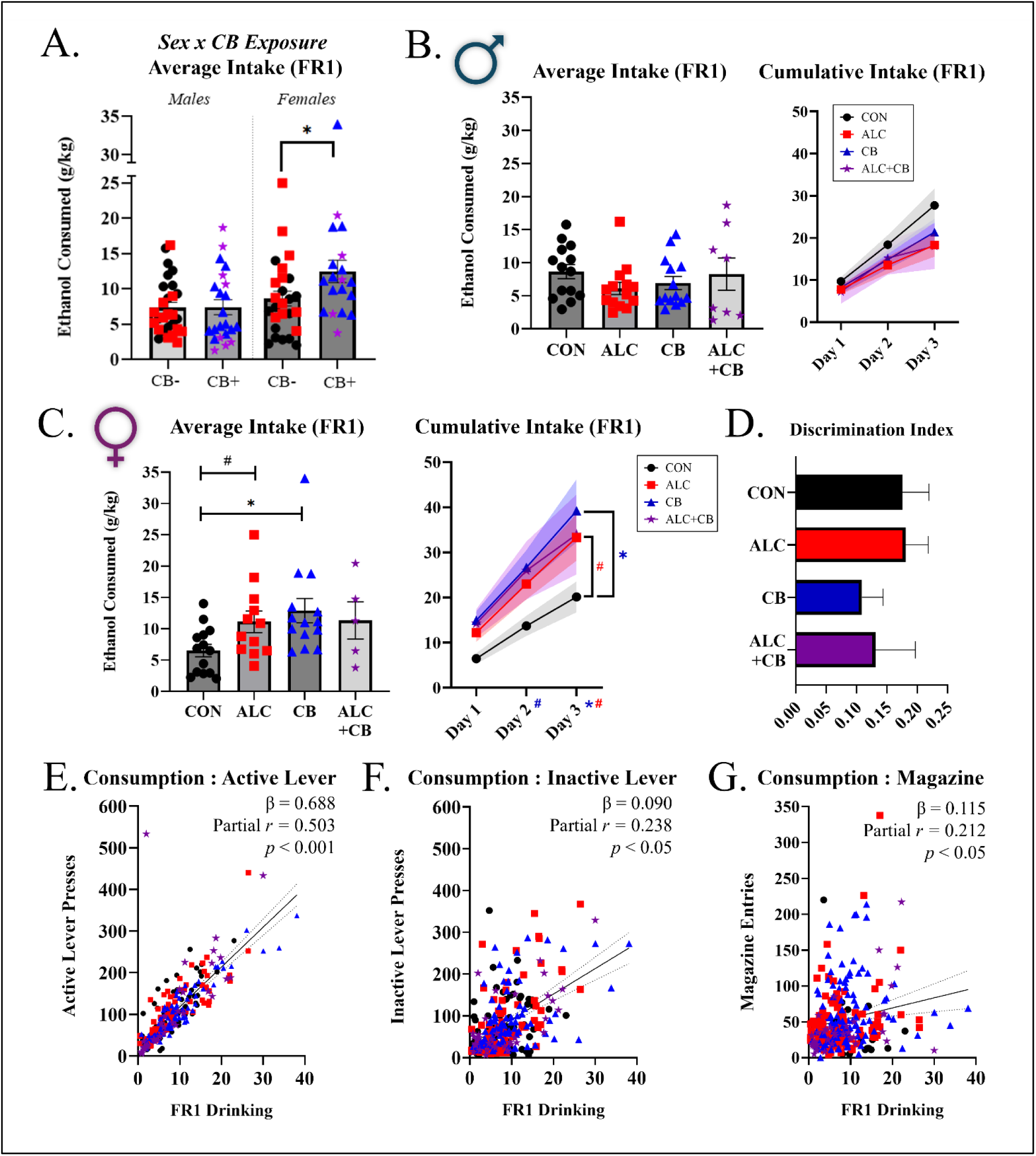
Ethanol consumption and operant-response measures during fixed-ratio 1 self-administration. (A) Mean ethanol intake across the three fixed-ratio 1 (FR1) sessions, separated by sex and prenatal cannabinoid exposure. CB− includes the CON and ALC groups, whereas CB+ includes the CB and ALC+CB groups. Female CB+ offspring consumed more ethanol than female CB− offspring. (B, C) Mean and cumulative ethanol intake across FR1 sessions in males (B) and females (C). No exposure-group differences were observed in males. In females, mean intake was greater in CB offspring than CON offspring and showed a trend toward greater intake in ALC offspring than CON offspring. For cumulative intake, CB offspring showed a trend toward greater intake than CON offspring on Day 2 and significantly greater intake on Day 3; ALC offspring showed a trend toward greater intake than CON offspring on Day 3. (D) Lever-discrimination index, calculated as (active lever presses − inactive lever presses)/(active lever presses + inactive lever presses). No significant effects of prenatal exposure were observed. (E–G) Associations between subject-level mean ethanol intake and mean active lever presses (E), inactive lever presses (F), and magazine entries (G) across available FR1 sessions. Active lever presses showed the strongest independent association with ethanol intake, with smaller independent associations observed for inactive lever presses. Partial correlations controlled for the other two operant-response measures. Standardized coefficients were obtained from a mixed-effects regression that additionally accounted for FR1 Day and repeated sessions within subjects. Solid lines in E–G show the unadjusted linear relationships between subject-level means; dotted lines indicate 95% confidence intervals. Bars and symbols represent mean ± SEM and individual subjects, respectively. Pairwise comparisons were Sidak-adjusted. Colored significance symbols in C indicate comparisons with CON for the corresponding exposure group. #*p* < .10; \**p* < .05

Active-lever responding also showed a Sex × ALC × CB interaction [*F*(1,86.52)=4.62, *p*=.034] (Table S10). Across offspring, discrimination favored the ethanol-associated lever and did not differ by sex or exposure (Fig. 3D). FR1 ethanol intake was correlated with active-lever presses [*r*=.596, *p* < .001] and inactive-lever presses [*r*=.503, *p*<.001], but not magazine entries [*r*=.173, *p*=.095] (Fig. 3E–G). In a repeated-session model, active-lever presses were the strongest unique correlate of intake (standardized β=.688, 95% CI [.606, .770], *p*<.001; Tables S10–S12). Descriptive statistics (mean ± SEM) for all FR1 operant-response measures by sex and prenatal exposure group are reported in Table S13.

Under a progressive-ratio schedule, ethanol intake increased with concentration and revealed a Sex × ALC × CB interaction [*F*(1,88.17)=7.03, *p*=.010] (Fig. 4; Table S14). In males, ALC and CB exposure interacted [*F*(1,46.05)=6.78, *p*=.012]; at 40% ethanol, ALC+CB offspring consumed more than CON, ALC-only, and CB-only offspring (all *ps* < .001; Table S15). In females, CB exposure interacted with concentration [*F*(2,60.80)=5.11, *p*=.009]; at 40%, all three exposed groups consumed more than CON (all *ps* ≤ .017; Table S16). Total liquid consumption varied with concentration and revealed concentration-dependent effects of ALC and CB exposure (all *ps* ≤ .025), as well as a Sex × ALC × CB interaction [*F*(1,87.97)=5.23, *p*=.025]. At 40%, ALC+CB males consumed more liquid than CB-only males (*p*=.032), whereas CB-only females consumed more than CON females (*p*=.040), with a trend toward greater consumption in ALC-only females (*p*=.080; Tables S17–S19). Breakpoint analyses likewise identified a Sex × ALC × CB interaction [*F*(1,88.12)=8.07, *p*=.006]. Male breakpoint showed an ALC × CB interaction (*p*=.017) without significant pairwise differences, whereas CB-only females reached a higher breakpoint than CON females at 40% ethanol (*p*=.047; Tables S20–S22).

**Figure 4.**
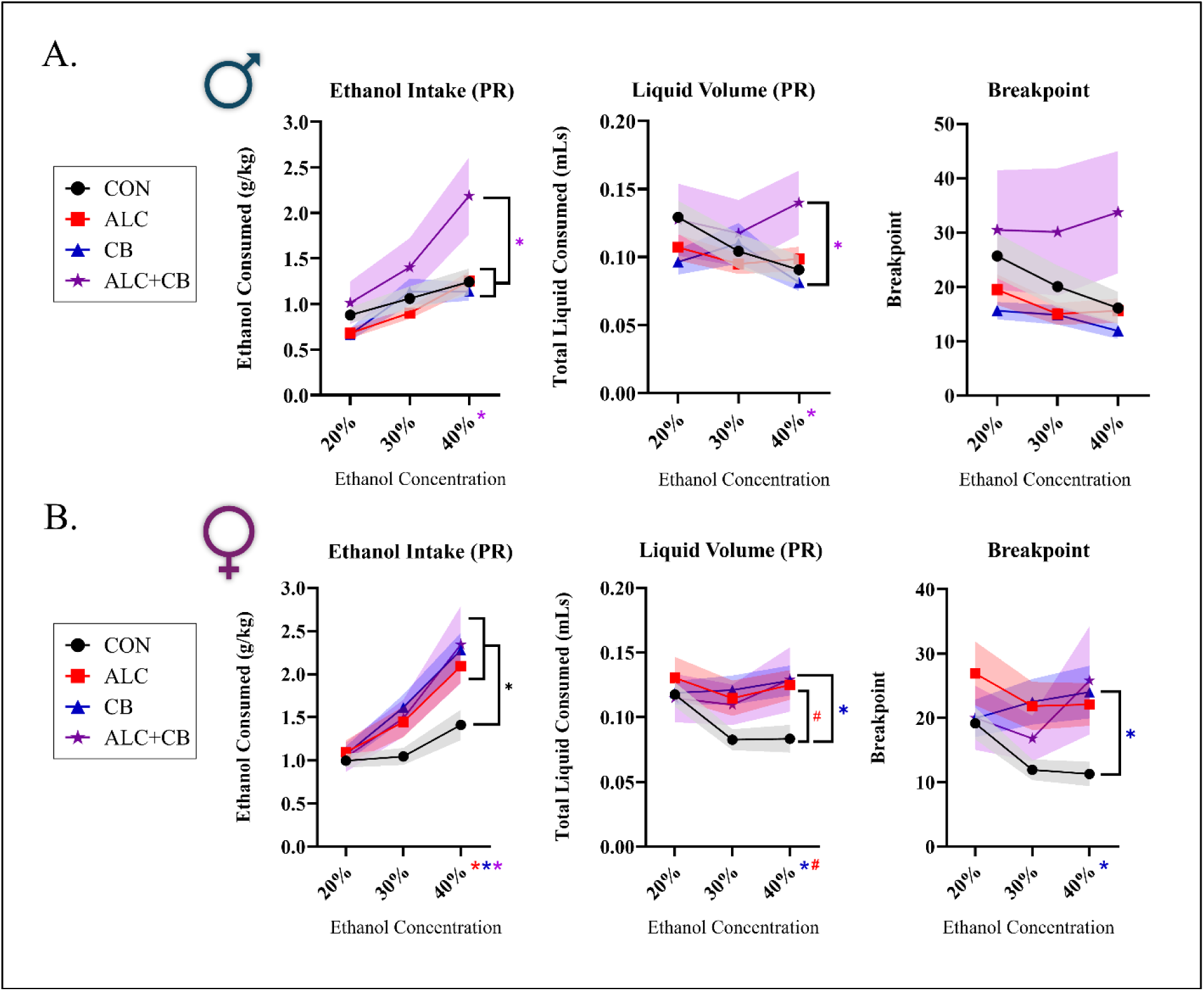
Prenatal alcohol and cannabinoid exposure alter progressive-ratio ethanol self-administration in a sex- and concentration-dependent manner. Ethanol intake (g/kg), total liquid consumed (mL), and breakpoint responding were assessed during progressive-ratio (PR) sessions using 20%, 30%, and 40% ethanol in (A) male and (B) female offspring. Lines and symbols represent means, and shaded regions represent SEM. In males, ALC+CB offspring consumed more ethanol than CON, ALC-only, and CB-only offspring at 40% ethanol, and more total liquid than CB-only offspring. No exposure-group differences in male breakpoint responding were observed at any concentration. In females, ALC-only, CB-only, and ALC+CB offspring consumed more ethanol than controls at 40%. At this concentration, CB-only offspring also consumed more total liquid and exhibited higher breakpoint responding than controls, while ALC-only offspring showed a trend toward greater liquid consumption. Data were analyzed using linear mixed-effects models with ethanol concentration as a repeated factor, followed by sex-stratified models and Sidak-adjusted comparisons within each concentration. Brackets denote pairwise comparisons. \**p* < .05; #*p* < .10.

### Prenatal ALC+CB exposure increased ethanol-seeking persistence during extinction and reinstatement in male offspring

Extinction responding revealed a Sex × ALC × CB interaction [*F*(1,89.96)=10.25, *p*=.002] (Fig. 5; Table S23). Sex-stratified analyses identified ALC × CB interactions for raw active-lever responding in males [*F*(1,46.80)=6.04, *p*=.018] and females [*F*(1,41.80)=7.33, *p*=.010]; significant pairwise differences were limited to greater responding in ALC+CB males compared to ALC-only and CB-only males (both *ps* ≤ .033; Fig. 5A,C; Tables S24–S25). When extinction responding was normalized to FR1 baseline, the Sex × ALC × CB interaction remained significant *[F*(1,85.89)=9.15, *p*=.003]. ALC+CB males showed greater normalized responding than CON, ALC-only, and CB-only males (all *ps* ≤ .0229; Tables S26–S28; Fig. 5B). CON, ALC-only, and CB-only males responded below baseline, whereas ALC+CB males did not (*p* = .597). No exposure-group differences in normalized extinction responding were detected in females (Fig. 5D).

**Figure 5.**
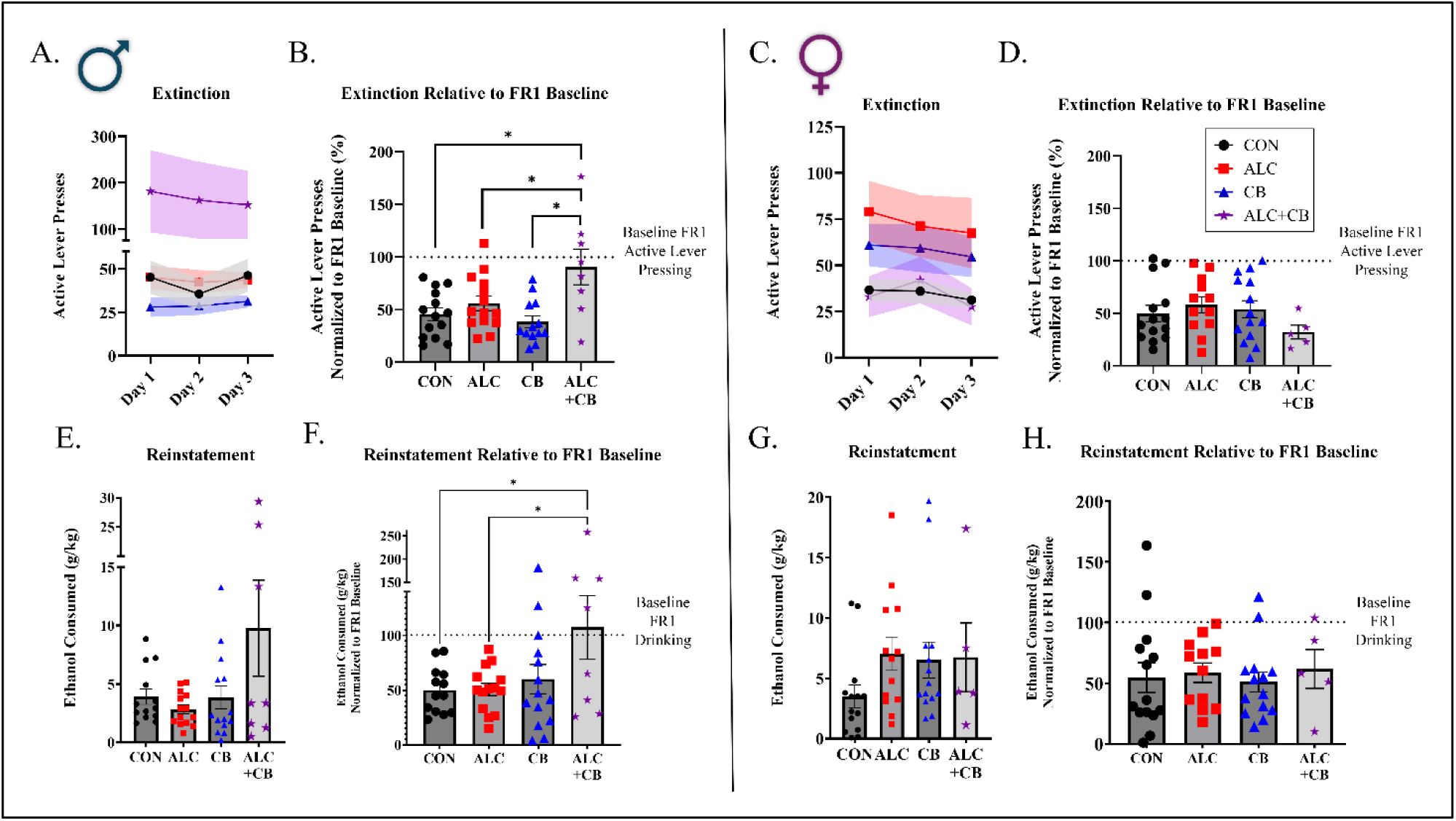
Prenatal ALC and CB exposures increased extinction and reinstatement ethanol-directed behavior selectively in male offspring. Active lever pressing across the three extinction sessions is shown for (A) male and (C) female offspring. Lines represent group means, and shaded regions represent the SEM. Extinction responding averaged across sessions was normalized to each subject’s mean active lever pressing during FR1 and is shown for (B) male and (D) female offspring. Ethanol intake during reinstatement is shown in g/kg for (E) male and (G) female offspring. Reinstatement intake normalized to each subject’s mean FR1 ethanol intake is shown for (F) male and (H) female offspring. Dotted lines at 100% indicate responding or intake equivalent to FR1 baseline. Bars represent means ± SEM, and individual symbols represent offspring. Brackets identify significant Šídák-adjusted exposure-group comparisons. * indicates *p* < .05. ALC, prenatal alcohol exposure; CB, prenatal cannabinoid exposure; CON, control; FR1, fixed-ratio 1.

Reinstatement ethanol intake also revealed a Sex × ALC × CB interaction [*F*(1,87)=5.55, *p*=.021] (Fig. 5E–H; Table S29). After normalization to FR1 intake, males showed a CB main effect [*F*(1,45)=6.44, *p*=.015], and ALC+CB males exhibited greater reinstatement intake relative to baseline than CON and ALC males (both *ps* =.039; Table S30). CON, ALC-only, and CB-only males reinstated below their FR1 baselines, whereas ALC+CB males did not (*p* = .808). No exposure effects were detected in females.

### Prenatal exposures produced sex- and region-specific changes in CNR1 and CNRIP1 expression

CNR1 expression was higher in the DMS than the mPFC [*F*(1,47.76)=20.54, *p*<.001], and showed CB × Region, ALC × CB × Region, and Sex × CB × Region interactions (all *ps* ≤ .019; Fig. 6A-B; Table S31). CNRIP1 expression was also higher in the DMS *[F*(1,47.48)=174.82, *p*<.001], and showed an ALC × CB interaction across sexes and regions *[F*(1,47.55)=6.34, *p*=.015] (Table S32).

**Figure 6.**
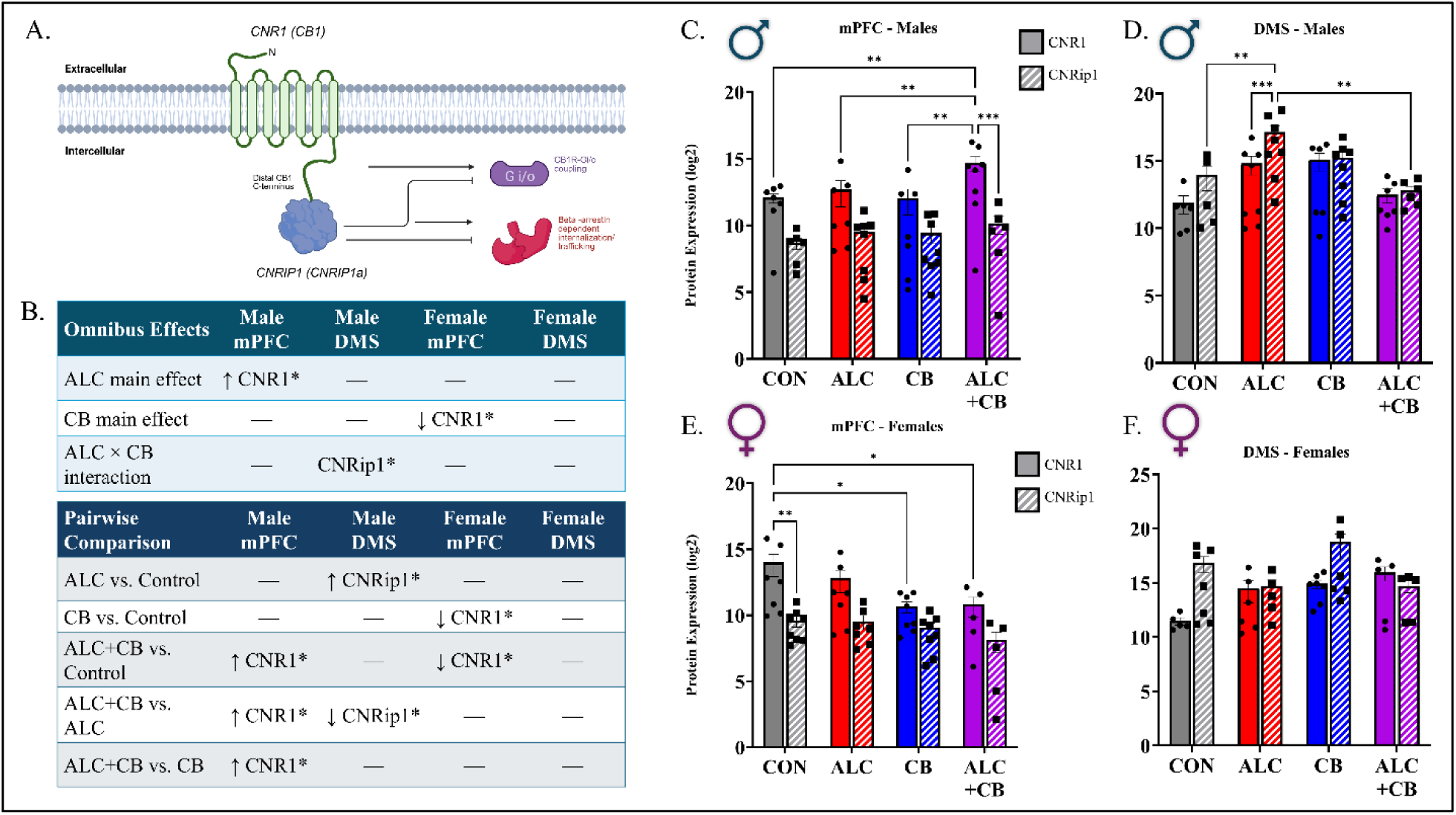
Prenatal ALC and CB exposures produced sex- and brain region-specific changes in CNR1 and CNRip1 protein expression. (A) Schematic depicting the cannabinoid receptor 1 (CNR1/CB1) and its interaction with cannabinoid receptor-interacting protein 1a (CNRip1/CNRIP1a), which can influence CB1 receptor signaling and trafficking. (B) Summary of significant exposure-related omnibus effects and selected exposure-group comparisons within each sex and brain region. Arrows indicate the direction of expression differences, and dashes indicate that no significant effect or selected comparison was detected. Log2-transformed CNR1 and CNRip1 protein expression is shown in the (C) male mPFC, (D) male DMS, (E) female mPFC, and (F) female DMS. Solid bars represent CNR1, and hatched bars represent CNRip1. Bars represent means ± SEM, and individual symbols represent offspring. CNR1 and CNRip1 were analyzed jointly within each sex and region using mixed-effects models with Protein as a matched factor and ALC and CB exposure as fixed factors. Brackets identify selected significant Šídák-adjusted comparisons. * indicates *p* < .05, ** indicates *p* < .01, and *** indicates *p* < .001.

Within the male mPFC, ALC+CB offspring had higher CNR1 expression than CON, ALC-only, and CB-only offspring (all Šídák-adjusted *ps* ≤ .010; Fig. 6C; Table S33). In the male DMS, ALC-only offspring had higher CNRIP1 expression than CON and ALC+CB offspring (all *ps* ≤ .016; Fig. 6D; Table S34). Within the female mPFC, CB-only and ALC+CB offspring had lower CNR1 expression than CON offspring (both *ps* ≤ .045; Fig. 6E; Table S35). No exposure-related differences were detected in the female DMS (Fig. 6F; Table S36).

### mPFC CNR1 expression was associated with ethanol-directed behaviors in male offspring

Among the four region-specific protein measures, mPFC CNR1 was the only measure associated with the combined behavioral profile of ethanol-seeking behaviors, and this association was detected exclusively in males [Pillai’s trace = .353, *F*(4,22)=3.01, *p*=.040, partial η²=.353] (Fig. 7A; Table S37). The omnibus association did not remain significant after Benjamini-Hochberg correction across the four protein models (*q*=.160).

**Figure 7.**
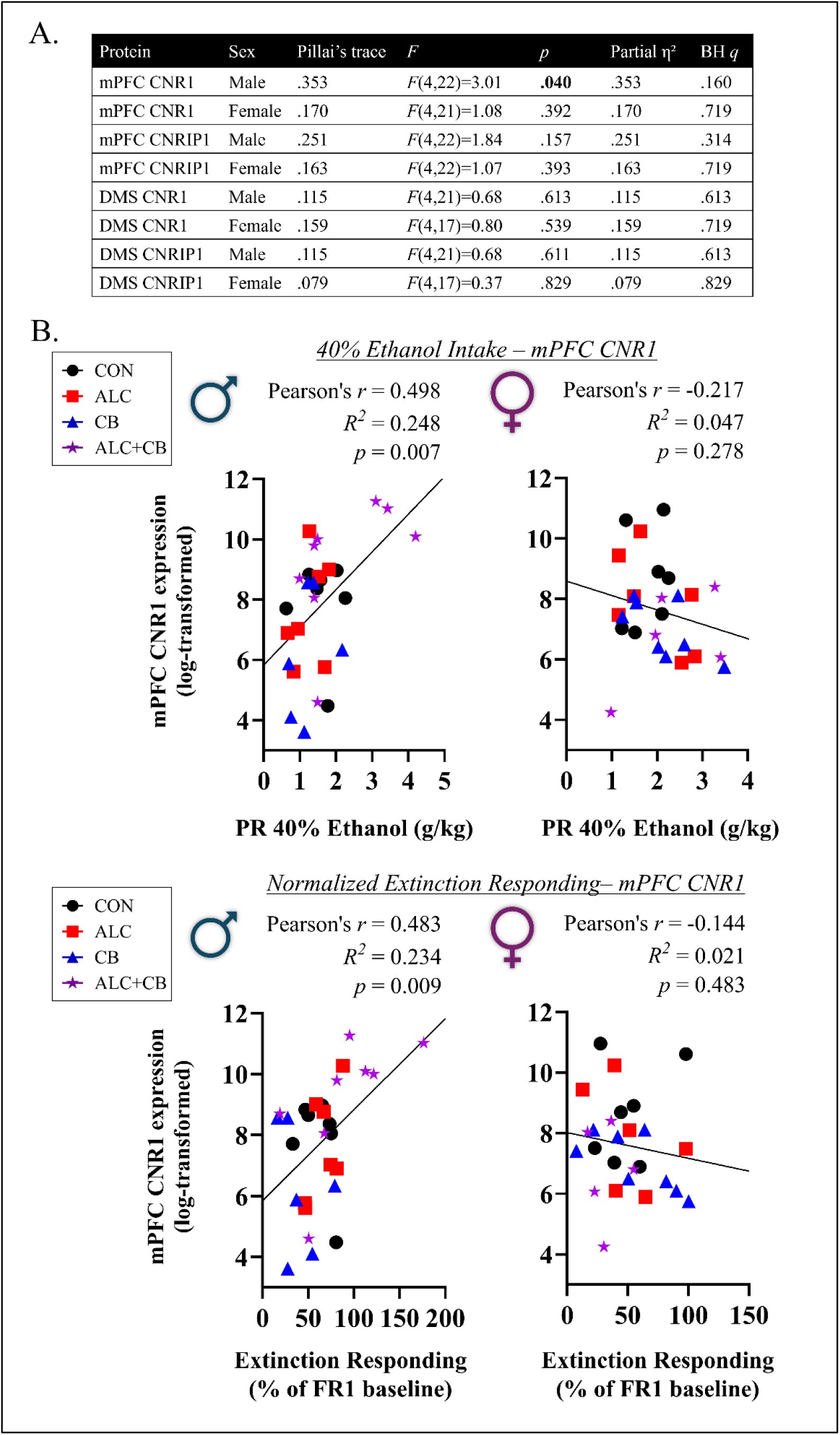
Associations between region-specific CNR1 and CNRIP1 expression and ethanol-directed behaviors. (A) Sex-stratified multivariate general linear models tested whether each region-specific protein measure was associated with a combined behavioral profile comprising average fixed-ratio 1 (FR1) ethanol intake, ethanol intake during the 40% progressive-ratio (PR) session, normalized extinction responding, and normalized reinstatement intake. Unadjusted *p* values and Benjamini-Hochberg (BH)-adjusted *q* values across the four protein measures within each sex are shown; bold indicates *p* < .05. (B) Pearson correlations between log-transformed mPFC CNR1 expression and ethanol intake during the 40% PR session (top) or extinction responding normalized to FR1 baseline (bottom), shown separately for males and females. mPFC, medial prefrontal cortex; DMS, dorsomedial striatum; BH, Benjamini-Hochberg.

In individual experiment follow-up analyses, higher mPFC CNR1 expression in males was positively correlated with FR1 ethanol intake, 40% progressive-ratio intake, normalized extinction responding, and normalized reinstatement intake [*r*s >.400, *ps* <.034] (Fig. 7B; Table S38). All four Pearson correlations remained significant after correction across the behavioral outcomes (*q*s<.034). Sensitivity analyses using Spearman correlations recapitulated the progressive-ratio, extinction, and reinstatement associations.

## Discussion

In this study, we found that prenatal alcohol and cannabinoid exposure during a critical window of cortical plate neurogenesis increased alcohol consumption in a sex and context-specific manner in adult offspring (Postnatal Day 200+). Prenatal cannabinoid exposure increased voluntary social ethanol intake, primarily in males, while combined exposure selectively augmented male ethanol consumption at the highest progressive-ratio concentration and increased responding during extinction and reinstatement. Notably, all three prenatal exposure conditions increased consumption of 40% ethanol in female offspring. Combined exposure also selectively increased medial prefrontal cortex CNR1 protein abundance in males, which was positively associated with ethanol intake and seeking. Thus, co-exposure produced a behavioral and molecular phenotype in mature adult offspring that was distinct from exposure to either substance individually, demonstrating that its consequences cannot be inferred from alcohol or cannabinoid exposure alone.

Importantly, exposure effects differed across sexes and testing conditions. During social home-cage access, prenatal cannabinoid exposure increased cumulative ethanol intake, primarily in males, whereas females consumed more ethanol than males overall, consistent with established sex differences in C57BL/6J mice [35, 36]. Prenatal exposure did not change low-demand fixed-ratio intake, but group differences emerged under a progressive-ratio schedule, particularly at the highest ethanol concentration. At 40% ethanol, all three exposed female groups consumed more than controls, and cannabinoid-only females reached a higher breakpoint. In males, combined exposure selectively increased 40% ethanol intake relative to control and both single-exposure groups, prolonged responding during extinction, and elevated reinstatement responding. Therefore, prenatal exposure produced different outcomes across sexes and behavioral contexts.

Prior studies have demonstrated that prenatal alcohol and cannabinoid exposure can increase later ethanol use, although reported outcomes vary by sex and task [7, 37–41]. Most directly, Ornelas et al. found that prenatal CP-55,940 exposure increased adolescent home-cage intake in both sexes and adult operant intake in females [24]. The present investigation extends these prenatal cannabinoid-associated increases in ethanol consumption beyond young adulthood, and further demonstrates that alcohol and cannabinoid co-exposure produces a distinct male profile of escalated ethanol-seeking.

Prenatal exposure affected motivation to work for and consume ethanol as concentration increased: specifically, control offspring reduced their intake, whereas exposed offspring maintained or increased intake at 40% ethanol. This divergence may reflect altered sensitivity to ethanol’s sensory or postabsorptive properties. PAE can attenuate aversion to ethanol’s bitter-like taste and enhance responsiveness to its odor in adolescent rats, although the taste effect is reported to dissipate by adulthood [42]. Prenatal exposure could also modify ethanol’s interoceptive or intoxicating effects or its pharmacokinetics. In this investigation, ethanol concentration increased sequentially, meaning that concentration was confounded with session order and accumulated operant experience. Thus, the present experiment cannot determine whether concentration, operant experience, or both drove this divergence. Future counterbalanced concentration testing with sensory and pharmacokinetic measures can help distinguish these mechanisms.

Prenatal alcohol and cannabinoid exposure also altered CNR1 and CNRIP1 protein abundance according to sex and brain region. In the male mPFC, factorial analysis identified an overall effect of PAE across CNR1 and CNRIP1 abundance, while pairwise comparisons localized group differences to CNR1, with combined exposure increasing CNR1 relative to control and both single-exposure groups. In the female mPFC, cannabinoid exposure reduced protein abundance overall, with lower CNR1 following prenatal cannabinoid and combined exposure compared to controls. In the male DMS, alcohol exposure alone increased CNRIP1 relative to control and combined exposure, whereas neither protein differed in the female DMS. CNRIP1 regulates CNR1 coupling, signaling, and internalization; therefore, increased DMS CNRIP1 may alter receptor regulation despite unchanged CNR1 abundance [43, 44]. Developmental alcohol and cannabinoid exposure has likewise produced sex-, region-, and model-dependent increases, reductions, or functional changes in CNR1 [45–48].

Combined exposure produced the clearest convergence between behavioral and protein outcomes. Among the protein-region pairings examined, only mPFC CNR1 was associated with the combined behavioral profile in males. Greater mPFC CNR1 abundance was positively associated with fixed-ratio intake, 40% progressive-ratio intake, extinction responding, and reinstatement responding. Pharmacological inhibition of CNR1 has been shown to reduce ethanol self-administration, habitual responding, and cue-induced reinstatement, whereas increasing endocannabinoid availability increases progressive-ratio responding [29, 49]. Alcohol can also strengthen glutamatergic transmission from the mPFC to direct-pathway DMS neurons, and manipulating these synapses bidirectionally alters alcohol seeking [32, 50]. The elevation of mPFC CNR1 and its positive behavioral associations therefore implicate altered cannabinoid regulation of corticostriatal signaling in the male co-exposure phenotype. However, as protein was measured in bulk tissue after behavioral testing, future experiments are required to identify the affected cells or projections, establish altered receptor function, or distinguish a prenatal exposure effect from adaptation to adult ethanol experience.

Our factorial exposure paradigm directly compared alcohol, cannabinoids, their combination, and controls within a single cohort, allowing effects unique to co-exposure to be distinguished from those produced by either substance alone. These effects emerged when ethanol access began around Postnatal Day 200, demonstrating that prior adolescent or young-adult ethanol experience was not required for prenatal exposure to increase later-adult consumption. However, several limitations should be acknowledged. First, home-cage intake was measured at the cage level and could not be assigned to individual offspring or related directly to individual operant and protein outcomes. Moreover, home-cage drinking occurred in social dyads, whereas operant testing occurred individually, preventing social context from being separated from drinking procedure. This distinction is important because social housing alters ethanol intake in mice [51, 52]. In follow-up investigations, radio-frequency identification-based systems can quantify individual ethanol intake while preserving social housing [52]. Our four-day prenatal exposure window intentionally targeted cortical neurogenesis [53], and this paradigm produced neurobehavioral differences in young adult offspring from this cohort [26]. However, exposure timing, dose, duration, route, and cannabinoid identity differ across developmental models and likely contribute to differences among studies [24, 37, 38, 41, 54]. Systematic comparisons that vary these parameters are needed to determine how each shapes the direction and duration of offspring outcomes. Finally, exposure-related morbidity previously reported in this cohort reduced the number of co-exposed females, limiting statistical power [26]. This imbalance is particularly important when interpreting the absence of co-exposure effects during fixed-ratio responding in females.

Taken together, the behavioral outcomes and exposure-dependent changes in CNR1 and CNRIP1 demonstrate that altered ethanol consumption and operant responding occurred alongside lasting changes within reward-related neural systems. In particular, the selective elevation of mPFC CNR1 in co-exposed males and its association with ethanol intake, extinction, and reinstatement identify cannabinoid signaling as a candidate contributor to later vulnerability to alcohol misuse. This behavioral-molecular convergence provides a rationale for testing whether interventions targeting cannabinoid signaling can attenuate or reverse exposure-induced increases in alcohol consumption and seeking. Our findings also support an incremental harm-reduction framework for prenatal polysubstance exposure; several outcomes occurred only after combined exposure, suggesting that eliminating either alcohol or cannabinoids from co-use may reduce those specific effects when complete cessation is not achievable. Together with the growing literature on prenatal polysubstance exposure, these data support comprehensive, nonjudgmental disclosure and documentation of all substances used during pregnancy so that exposure combinations can inform monitoring of exposure-specific outcomes. The observed sex differences further support consideration of both exposure history and biological sex in clinical phenotyping, diagnostic assessment, and longitudinal follow-up.

## Acknowledgments

Research reported in this publication was supported by the National Institute on Alcohol Abuse and Alcoholism of the National Institutes of Health (F32AA029866 and K99AA032336 to SKR, R01AA028406 to RCM), and by the National Institute of General Medical Sciences (K12GM154716).

## CRediT Author Statement

Siara K. Rouzer: Conceptualization, Methodology, Formal Analysis, Investigation, Data Curation, Writing - Original Draft, Writing - Review & Editing, Visualization, Project Administration, Funding Acquisition. Abigail Bowring: Investigation, Data Curation, Formal Analysis, Writing - Original Draft, Visualization. Aisley George: Investigation, Data Curation, Writing - Original Draft. McKay Domen: Investigation, Data Curation. Emma Labbe: Investigation, Data Curation. Marisol Palacios: Investigation, Data Curation. Rajesh C. Miranda: Writing - Review & Editing, Funding Acquisition.

Figures 1 and 6, as well as the graphical abstract, were created using BioRender.

## References

1. Gowing, L.R., et al., Global statistics on addictive behaviours: 2014 status report. Addiction, 2015. 110(6): p. 904–919.

2. Rehm, J., et al., Global burden of disease and injury and economic cost attributable to alcohol use and alcohol-use disorders. The lancet, 2009. 373(9682): p. 2223–2233.

3. England, L.J., Alcohol use and co-use of other substances among pregnant females aged 12–44 years—United States, 2015–2018. MMWR. Morbidity and Mortality Weekly Report, 2020. 69.

4. Verhulst, B., M.C. Neale, and K.S. Kendler, The heritability of alcohol use disorders: a meta-analysis of twin and adoption studies. Psychol Med, 2015. 45(5): p. 1061–72.

5. Kranzler, H.R., et al., Genome-wide association study of alcohol consumption and use disorder in 274,424 individuals from multiple populations. Nature Communications, 2019. 10(1): p. 1499.

6. Alati, R., et al., In utero alcohol exposure and prediction of alcohol disorders in early adulthood: a birth cohort study. Archives of general psychiatry, 2006. 63(9): p. 1009–1016.

7. Baer, J.S., et al., A 21-year longitudinal analysis of the effects of prenatal alcohol exposure on young adult drinking. Archives of general psychiatry, 2003. 60(4): p. 377–385.

8. Goldschmidt, L., et al., Prenatal alcohol exposure and offspring alcohol use and misuse at 22 years of age: a prospective longitudinal study. Neurotoxicology and teratology, 2019. 71: p. 1–5.

9. Fabio, M.C., et al., Prenatal ethanol increases ethanol intake throughout adolescence, alters ethanol-mediated aversive learning, and affects μ but not δ or κ opioid receptor mRNA expression. European Journal of Neuroscience, 2015. 41(12): p. 1569–1579.

10. Fabio, M.C., et al., Prenatal ethanol exposure increases ethanol intake and reduces c-Fos expression in infralimbic cortex of adolescent rats. Pharmacology Biochemistry and Behavior, 2013. 103(4): p. 842–852.

11. Gore-Langton, J.K. and L.P. Spear, Prenatal ethanol exposure attenuates sensitivity to the aversive effects of ethanol in adolescence and increases adult preference for a 5% ethanol solution in males, but not females. alcohol, 2019. 79: p. 59–69.

12. Randall, C.L., et al., Effect of prenatal alcohol exposure on consumption of alcohol and alcohol-induced sleep time in mice. Pharmacology Biochemistry and Behavior, 1983. 18: p. 325–329.

13. Grant, B.F., et al., Epidemiology of DSM-5 Alcohol Use Disorder: Results From the National Epidemiologic Survey on Alcohol and Related Conditions III. JAMA Psychiatry, 2015. 72(8): p. 757–766.

14. Rouzer, S.K., et al., Alcohol & cannabinoid co-use: Implications for impaired fetal brain development following gestational exposure. Experimental neurology, 2023. 361: p. 114318.

15. Subbaraman, M.S. and W.C. Kerr, Simultaneous versus concurrent use of alcohol and cannabis in the National Alcohol Survey. Alcoholism: clinical and experimental research, 2015. 39(5): p. 872–879.

16. Eisenberg, L. and S.S. Brown, The best intentions: Unintended pregnancy and the well-being of children and families. 1995.

17. Metrik, J., et al., Sexual risk behavior and heavy drinking among weekly marijuana users. Journal of studies on alcohol and drugs, 2016. 77(1): p. 104–112.

18. Simons, J.S., S.A. Maisto, and T.B. Wray, Sexual risk taking among young adult dual alcohol and marijuana users. Addictive Behaviors, 2010. 35(5): p. 533–536.

19. Green, K.M., et al., Outcomes associated with adolescent marijuana and alcohol use among urban young adults: A prospective study. Addictive behaviors, 2016. 53: p. 155–160.

20. Magill, M., et al., The role of marijuana use in brief motivational intervention with young adult drinkers treated in an emergency department. Journal of studies on alcohol and drugs, 2009. 70(3): p. 409–413.

21. Abel, E. and M. Subramanian, Effects of low doses of alcohol on delta-9-tetrahydrocannabinol’s effects in pregnant rats. Life sciences, 1990. 47(18): p. 1677–1682.

22. Hartman, R.L., et al., Controlled cannabis vaporizer administration: blood and plasma cannabinoids with and without alcohol. Clinical chemistry, 2015. 61(6): p. 850–869.

23. Breit, K.R., et al., Combined vapor exposure to THC and alcohol in pregnant rats: maternal outcomes and pharmacokinetic effects. Neurotoxicology and teratology, 2020. 82: p. 106930.

24. Ornelas, L.C., et al., The impact of prenatal alcohol, synthetic cannabinoid and co-exposure on behavioral adaptations in adolescent offspring and alcohol self-administration in adulthood. Neurotoxicology and Teratology, 2024. 102: p. 107341.

25. Abel, E.L., Alcohol enhancement of marihuana-induced fetotoxicity. Teratology, 1985. 31(1): p. 35–40.

26. Rouzer, S.K., et al., Early life outcomes of prenatal exposure to alcohol and synthetic cannabinoids in mice. Drug and Alcohol Dependence Reports, 2025. 16: p. 100356.

27. Breit, K.R., et al., Effects of prenatal alcohol and delta-9-tetrahydrocannabinol exposure via electronic cigarettes on motor development. Alcoholism: clinical and experimental research, 2022. 46(8): p. 1408–1422.

28. Rouzer, S.K., A. Sreeram, and R.C. Miranda, Reduced fetal cerebral blood flow predicts perinatal mortality in a mouse model of prenatal alcohol and cannabinoid exposure. BMC Pregnancy and Childbirth, 2024. 24(1): p. 263.

29. Economidou, D., et al., Role of cannabinoidergic mechanisms in ethanol self-administration and ethanol seeking in rat adult offspring following perinatal exposure to Delta9-tetrahydrocannabinol. Toxicol Appl Pharmacol, 2007. 223(1): p. 73–85.

30. Hausknecht, K., et al., Prenatal ethanol exposure persistently alters endocannabinoid signaling and endocannabinoid-mediated excitatory synaptic plasticity in ventral tegmental area dopamine neurons. The Journal of Neuroscience, 2017. 37(24): p. 5798–5808.

31. Ruiz-Calvo, A., et al., Pathway-Specific Control of Striatal Neuron Vulnerability by Corticostriatal Cannabinoid CB1 Receptors. Cereb Cortex, 2018. 28(1): p. 307–322.

32. Xie, X., et al., Linking input- and cell-type-specific synaptic plasticity to the reinforcement of alcohol-seeking behavior. Neuropharmacology, 2023. 237: p. 109619.

33. Smith, T.H., et al., Cannabinoid receptor-interacting protein 1a modulates CB1 receptor signaling and regulation. Mol Pharmacol, 2015. 87(4): p. 747–65.

34. Blegen, M.B., et al., Alcohol operant self-administration: Investigating how alcohol-seeking behaviors predict drinking in mice using two operant approaches. Alcohol, 2018. 67: p. 23–36.

35. Rivera-Irizarry, J.K., et al., Sex differences in binge alcohol drinking and the behavioral consequences of protracted abstinence in C57BL/6J mice. Biology of sex differences, 2023. 14(1): p. 83.

36. Sneddon, E.A., R.D. White, and A.K. Radke, Sex differences in binge-like and aversion-resistant alcohol drinking in C57 BL/6J mice. Alcoholism: clinical and experimental research, 2019. 43(2): p. 243–249.

37. Brancato, A., et al., In utero Δ9-tetrahydrocannabinol exposure confers vulnerability towards cognitive impairments and alcohol drinking in the adolescent offspring: Is there a role for neuropeptide Y? Journal of Psychopharmacology, 2020. 34(6): p. 663–679.

38. Castelli, V., et al., Prenatal THC exposure and binge-like alcohol drinking in early adolescence: From sex-specific drinking vulnerability to abnormal endocannabinoid-dopamine nexus in the nucleus accumbens. J Psychopharmacol, 2026. 40(9): p. 1493–1505.

39. Chotro, M.G., C. Arias, and G. Laviola, Increased ethanol intake after prenatal ethanol exposure: studies with animals. Neuroscience & Biobehavioral Reviews, 2007. 31(2): p. 181–191.

40. Duko, B., et al., Prenatal alcohol exposure and offspring subsequent alcohol use: A systematic review. Drug and Alcohol Dependence, 2022. 232: p. 109324.

41. Navarro, D., et al., Fetal cannabinoid syndrome: behavioral and brain alterations of the offspring exposed to dronabinol during gestation and lactation. International Journal of Molecular Sciences, 2024. 25(13): p. 7453.

42. Youngentob, S.L. and J.I. Glendinning, Fetal ethanol exposure increases ethanol intake by making it smell and taste better. Proc Natl Acad Sci U S A, 2009. 106(13): p. 5359–64.

43. Blume, L.C., et al., Cannabinoid receptor interacting protein (CRIP1a) attenuates CB1R signaling in neuronal cells. Cell Signal, 2015. 27(3): p. 716–726.

44. Booth, W.T., et al., Cannabinoid Receptor Interacting Protein 1a (CRIP1a): Function and Structure. Molecules, 2019. 24(20).

45. Bara, A., et al., Sex-dependent effects of in utero cannabinoid exposure on cortical function. Elife, 2018. 7.

46. Benevenuto, S.G.M., et al., Prenatal exposure to Cannabis smoke induces early and lasting damage to the brain. Neurochem Int, 2022. 160: p. 105406.

47. Di Bartolomeo, M., et al., Crosstalk between the transcriptional regulation of dopamine D2 and cannabinoid CB1 receptors in schizophrenia: Analyses in patients and in perinatal Delta9-tetrahydrocannabinol-exposed rats. Pharmacol Res, 2021. 164: p. 105357.

48. Oubraim, S., et al., Prenatal ethanol exposure causes anxiety-like phenotype and alters synaptic nitric oxide and endocannabinoid signaling in dorsal raphe nucleus of adult male rats. Transl Psychiatry, 2022. 12(1): p. 440.

49. Gianessi, C.A., et al., Endocannabinoid contributions to alcohol habits and motivation: Relevance to treatment. Addict Biol, 2020. 25(3): p. e12768.

50. Ma, T., et al., Bidirectional and long-lasting control of alcohol-seeking behavior by corticostriatal LTP and LTD. Nat Neurosci, 2018. 21(3): p. 373–383.

51. Fulenwider, H.D., et al., Social Housing Leads to Increased Ethanol Intake in Male Mice Housed in Environmentally Enriched Cages. Front Behav Neurosci, 2021. 15: p. 695409.

52. Petersen, N., et al., A novel mouse home cage lickometer system reveals sex-and housing-based influences on alcohol drinking. eneuro, 2024. 11(10).

53. Chen, V.S., et al., Histology Atlas of the Developing Prenatal and Postnatal Mouse Central Nervous System, with Emphasis on Prenatal Days E7.5 to E18.5. Toxicol Pathol, 2017. 45(6): p. 705–744.

54. Kokhan, V.S., et al., Sex-related differences in voluntary alcohol intake and mRNA coding for synucleins in the brain of adult rats prenatally exposed to alcohol. Biomedicines, 2022. 10(9): p. 2163.

