## Supplemental Materials for "Prenatal alcohol and cannabinoid co-exposure increases ethanol-seeking in adult offspring and produces sex-dependent medial prefrontal cannabinoid receptor 1 expression"

### Supplementary Methods

#### Experimental animals and study design

Adult male and female C57BL/6J offspring were selected from our previously characterized prenatal exposure cohort [1], where breeding and prenatal exposure procedures have been reported previously. Briefly, pregnant dams were randomly assigned to control (CON), alcohol (ALC), cannabinoid (CB), or combined alcohol and cannabinoid (ALC+CB) conditions. From gestational days 12–15, dams received the synthetic CB1/CB2 receptor agonist CP-55,940 (0.75 mg/kg, i.p.) or vehicle immediately before a 30-min exposure to ethanol vapor or room air. The present cohort represented 22 litters, including eight CON, five ALC, six CB, and three ALC+CB litters. All offspring had previously completed open-field testing, and all but one ALC female completed rotarod testing [1]. The female that did not meet the rotarod habituation criterion remained eligible for open-field testing and the subsequent assessments reported here.

Animals were maintained on a 12:12-h light-dark cycle, with lights on at 07:00, and with access to food and water ad libitum. Animals were housed 2–3 per cage with offspring of the same sex and prenatal exposure condition. Single housing was used only when required to address cage-mate aggression in accordance with animal-welfare considerations. All experimental procedures were approved by the Texas A&M University Institutional Animal Care and Use Committee (Protocol No. 2021-0331).

Offspring completed the present assessments in a fixed sequence consisting of home-cage ethanol drinking, operant acquisition and ethanol fixed-ratio self-administration, progressive-ratio testing, extinction, reinstatement, and tissue collection. Home-cage drinking began at a mean postnatal age of 199.5 ± 5.8 days (6.6 ± 0.2 months) and ended approximately 19 days later. Operant training began at postnatal day 267.4 ± 5.1 (8.8 ± 0.2 months). Because subjects required between 5 and 15 sessions to meet the operant-acquisition criteria, the age at completion of operant testing varied among subjects. Tissue was collected at postnatal day 363.4 ± 11.6 (11.9 ± 0.4 months). The mean within-subject interval between completion of home-cage drinking and initiation of operant training was 49.9 ± 3.7 days, and the mean interval between completion of operant testing and tissue collection was 84.2 ± 8.4 days. Starting age and inter-assessment intervals did not differ by sex or prenatal exposure.

Home-cage drinking was assessed in 108 offspring housed across 46 cages. This cohort included 36 CON offspring (18 males, 18 females; 15 cages), 29 ALC offspring (14 males, 15 females; 13 cages), 30 CB offspring (14 males, 16 females; 13 cages), and 13 ALC+CB offspring (8 males, 5 females; 5 cages). All occupants contributed to the home-cage assessment, and the cage served as the experimental unit. Subsequent operant assessments included 96 offspring from all 22 litters: 28 CON (14 males, 14 females), 27 ALC (14 males, 13 females), 28 CB (14 males, 14 females), and 13 ALC+CB offspring (8 males, 5 females). To reduce overrepresentation of individual litters contributing four or more offspring of the same sex and exposure condition, animals were selected to maximize representation across litters and home cages. Two additional animals failed to meet the minimum response criterion of 100 total lever presses per night across three training nights and were excluded from operant analyses.

Protein measurements were obtained from 57 offspring representing all 22 litters, including 15 CON (7 males, 8 females), 15 ALC (8 males, 7 females), 16 CB (8 males, 8 females), and 11 ALC+CB offspring (6 males, 5 females). Analysis-specific sample sizes were reduced when measurements did not meet the protein data-quality or outlier criteria described below.

#### Voluntary home-cage ethanol drinking

Voluntary ethanol drinking was assessed in age-matched cohorts tested across 17 months. Identical procedures were used across cohorts, and the order in which cages were assessed was randomized. Drinking sessions were conducted Monday through Friday for three consecutive weeks, producing 15 drinking sessions over 19 calendar days. Each session lasted 3 h and began between 08:00 and 09:00 during the light phase.

Drinking tubes were constructed from 15-mL conical tubes fitted with Hydropac VX13 valves (HYP-5300-05; Hydropac LLC, Belcamp, MD) and secured using custom 3D-printed holders adapted from an open-source home-cage sipper design [2]. Holders were attached to the food tray at the center of each cage and secured with binder clips. Tubes were filled with approximately 11–13 mL of fluid and primed before placement by depressing the valve until fluid was dispensed. Each numbered tube was assigned to a specific cage and weighed with its cap secured immediately before the session. After placement, the cap was loosened to permit fluid dispensing while limiting evaporation and spillage.

A 20% ethanol solution was prepared by diluting 95% ethanol stock with water. During sessions 1–5, mice received access to one tube containing 20% ethanol. During sessions 6–15, a second tube containing water was added to assess ethanol-to-water preference. The left or right position of the ethanol tube was counterbalanced across cages but remained constant within each cage throughout testing. Standard cage water bottles were made inaccessible during the 3-h session and restored immediately afterward. At session completion, tube caps were secured before removal, and each tube was reweighed. No separate leakage or evaporation-control tubes were used. Measurements from sessions with observed tube leakage or failure of fluid dispensing were excluded.

Animals were weighed on the first day of home-cage testing. Daily cage-level ethanol intake was estimated from the decrease in ethanol-tube weight according to the following calculation: ethanol intake (g/kg) = [(pre-session tube weight [g] − post-session tube weight [g]) × 0.20] ÷ average body weight of cage occupants (kg).

Cumulative ethanol intake was calculated by summing daily cage-level intake across the 15 sessions. For preference calculations, decreases in ethanol- and water-tube weights were converted to fluid volumes using fixed conversion values of 0.95 g/mL for the ethanol solution and 1.00 g/mL for water. An ethanol-to-water ratio was calculated by dividing cumulative ethanol-solution volume by cumulative water volume across sessions 6–15. This ratio was natural-log transformed, such that 0 represented equal consumption of ethanol solution and water, positive values represented proportionally greater ethanol consumption, and negative values represented proportionally greater water consumption. Confirmed zero-consumption measurements were retained when tube function had been verified. Because consumption was measured collectively, the cage served as the experimental unit.

#### Operant ethanol self-administration, extinction, and reinstatement

Operant testing was conducted using a lever-based modification of the CUP paradigm [3]. Extra-wide modular mouse chambers (ENV-307W-CT; Med Associates, Fairfax, VT) were enclosed within sound-attenuating cubicles (ENV-022MD) and controlled using MED-PC software. Each chamber contained two retractable ultrasensitive levers (ENV-312-3W) mounted on the same wall, a stimulus light above the active lever, a chamber houselight, and a liquid receptacle equipped with a head-entry detector (ENV-303LPHDW) on the opposite wall. The right lever was designated as active for all animals, whereas the left lever was inactive. Ethanol was delivered from a syringe pump (PHM-107) through 18-gauge tubing into the liquid receptacle. Active- and inactive-lever presses, reward deliveries, and magazine entries were recorded throughout each session.

An even layer of familiar home-cage bedding was placed in the tray beneath the grid floor. Five pieces of standard chow were provided during overnight training, and three pieces were provided during daytime sessions. Water was not available inside the chamber. Chambers, grid floors, trays, and liquid receptacles were cleaned and dried between sessions.

Operant acquisition (magazine training) consisted of 12-h overnight FR1 sessions conducted Sunday through Thursday nights. Both levers were extended throughout each session. Each active-lever press delivered 18 µL of 20% ethanol into the liquid receptacle and illuminated the stimulus light above the active lever. Each delivery was followed by a 5-s timeout during which additional active-lever presses were recorded but did not produce another reward. Inactive-lever presses had no programmed consequence. Acquisition required at least 100 total lever presses and at least 70% active-lever responding during each of three consecutive sessions. Training continued until both criteria were met or for a maximum of 15 sessions. Animals that did not meet the criteria after 15 sessions proceeded to daytime testing.

Following acquisition, mice completed three FR1 sessions, three progressive-ratio sessions, three extinction sessions, and one reinstatement session. All daytime sessions lasted 3 h and were conducted on consecutive days, including weekends. During FR1 sessions, each active-lever press produced 18 µL of 20% ethanol and illumination of the stimulus light, followed by the 5-s timeout.

Progressive-ratio sessions were conducted using ethanol concentrations of 20%, 30%, and 40% on the first, second, and third sessions, respectively. The response requirement increased after each earned reward according to the programmed sequence 1, 2, 4, 9, 12, 15, 20, 25, 32, 40, 50, 62, 77, 95, 118, 145, 178, 219, 268, 328, 402, 492, 603, 737, 901, 1102, 1347, 1647, and 2012 active-lever presses. Completion of each response requirement produced an 18-µL ethanol reward, illumination of the stimulus light, and a 5-s timeout. Sessions continued for the full 3 h without an inactivity-based termination criterion. Breakpoint was defined as the final response requirement completed during the session.

During extinction, both levers remained available, but lever presses produced neither ethanol nor illumination of the stimulus light. After three extinction sessions, mice completed one ethanol-reinforced reinstatement session. Reinstatement was conducted under the same FR1 contingency used previously, with each active-lever press producing 18 µL of 20% ethanol and illumination of the stimulus light, followed by the 5-s timeout. No ethanol prime was placed in the receptacle before reinstatement.

Animals were weighed immediately before overnight acquisition (magazine) training. Syringe volumes were recorded immediately before and after each session. Any ethanol remaining in the liquid receptacle was recovered using a small syringe and measured. Consumed ethanol-solution volume was calculated as: consumed solution volume (mL) = starting syringe volume (mL) − ending syringe volume (mL) − recovered cup volume (mL). Ethanol intake was calculated as: ethanol intake (g/kg) = [consumed solution volume (mL) × ethanol concentration] ÷ body weight (kg).

Syringe-based measurements were used as the primary measure of consumption, whereas programmed reward deliveries were retained as a behavioral measure and quality-control reference. When the liquid receptacle flooded, no consumption value was assigned for that session. Lever presses, reward deliveries, and magazine entries from flooded sessions were retained and remained subject to the outcome-specific outlier criteria described below.

#### Blood ethanol collection following ethanol consumption assessments

As a planned component of the ethanol consumption assessments, 20 µL of tail blood was collected into heparinized capillary tubes immediately following the final home-cage drinking session, the final FR1 session, and the reinstatement session for measurement of blood ethanol concentrations (BECs) using gas chromatography. However, following analysis, ethanol was only detected in 30.3% of samples obtained after home-cage drinking, 48.1% of samples obtained after FR1 testing, and 34.0% of samples obtained after reinstatement. These low and variable detection rates are not considered reliable measures of ethanol consumption, particularly because operant cup-volume measurements confirmed ethanol consumption even when no ethanol was detected in the corresponding blood sample. C57BL/6J mice exhibit a relatively rapid rise and decline in BEC following ethanol administration [4]. Because subjects could consume ethanol at any point during each 3-h session, post-session BECs may have reflected the interval between the most recent ethanol consumption and blood collection more strongly than the total amount consumed. BECs are therefore not reported as outcomes of these assessments. However, all subjects underwent tail-blood collection at the same three testing points to control for potential stress-related effects of the collection procedure.

#### Tissue collection and Western blot analyses

Animals received an intraperitoneal injection of a ketamine/xylazine solution prepared in phosphate-buffered saline. The working solution contained 2.5 mg/mL ketamine hydrochloride (100 mg/mL stock; Dechra, Northwich, UK) and 10 mg/mL xylazine (100 mg/mL stock; Covetrus, Portland, ME, USA) and was administered at 0.1 mL/10 g body weight. Loss of the righting reflex and absence of a response to foot pinch were confirmed before secondary decapitation. Animals were not perfused.

Brains were removed immediately after decapitation, and the bilateral medial prefrontal cortex and dorsomedial striatum were manually dissected from fresh tissue. Regions were identified under a dissecting microscope with reference to the Allen Mouse Brain Atlas. Tissue was maintained on dry ice during dissection and weighing. Samples were homogenized by sonication in 1× RIPA buffer supplemented with Halt protease inhibitor at a ratio of 20 µL buffer per milligram of tissue. Homogenates were centrifuged at 20,800 × g for 10 min at 4°C. The resulting supernatant was transferred into aliquots and stored at −80°C until analysis. Samples were thawed on ice before use. Protein concentrations were determined using the Pierce BCA Protein Assay Kit (Thermo Fisher Scientific, catalog no. 23250) against a bovine serum albumin standard curve, with absorbance measured at 562 nm according to the manufacturer’s instructions. Each sample was prepared with 2 µg of protein in a total volume of 15 µL using 4× LI-COR loading buffer containing 10% β-mercaptoethanol, diluted to a final 1× loading-buffer concentration, and heated at 95°C for 5 min. The full 15-µL sample was loaded into each lane. Proteins were separated on 12% NuPAGE Bis-Tris gels in the 1.0-mm, 17-well format using 1× MOPS running buffer. Gels were run at 200 V until the dye front reached the bottom of the gel, approximately 42–45 min. Proteins were transferred to PVDF membranes using an Invitrogen iBlot Gel Transfer Stack and iBlot transfer system.

Following transfer, total protein was visualized on every membrane using REVERT Total Protein Stain (LI-COR Biosciences). Membranes were treated with methanol for 30 s, rinsed in Milli-Q water for 2 min, incubated in REVERT stain for 5 min, washed twice for 10 min in REVERT wash solution, rinsed three times in Milli-Q water, and imaged using an Odyssey CLx imaging system (LI-COR Biosciences). Membranes were subsequently blocked in LI-COR Blocking Buffer for 1 h at room temperature. Membranes were co-incubated overnight at 4°C with rabbit monoclonal anti-CNR1 antibody (clone EPR23934-20; Abcam, catalog no. ab259323; 1:100) and rabbit polyclonal anti-CNRIP1 antibody (Proteintech, catalog no. 16827-1-AP; 1:100) in LI-COR Blocking Buffer. Membranes were washed three times for 5 min in Tris-buffered saline containing approximately 0.05% Tween-20 (TBST), once for 3 min in TBS, and rinsed in Milli-Q water. Membranes were then incubated for 1 h at room temperature, protected from light, with IRDye 800CW goat anti-rabbit IgG secondary antibody (LI-COR Biosciences; 1:14,300) in LI-COR Blocking Buffer containing 0.01% SDS and 0.2% Tween-20. Following secondary incubation, membranes were washed three times for 5 min in TBST, once for 3 min in TBS, and rinsed in Milli-Q water.

Fluorescent signals were acquired using the Odyssey CLx and analyzed in Image Studio software (LI-COR Biosciences; version 5.2). CNR1 and CNRIP1 immunoreactivity was quantified at approximately 53 kDa and 20 kDa, respectively, using the Chameleon Duo Pre-stained Protein Ladder as a molecular-weight reference. Bands were manually identified using rectangular regions of interest. Double bands were not observed for either target. When smearing occurred, the entire smear was included in the quantified region. For within-membrane normalization, a lane normalization factor was calculated by dividing the REVERT total-protein signal for each lane by the highest total-protein lane signal on that membrane: lane normalization factor = total-protein signal in lane X ÷ highest total-protein lane signal on the membrane. The target-protein signal for each lane was then divided by its corresponding lane normalization factor: within-membrane normalized signal = target-protein signal ÷ lane normalization factor. To account for variation among membranes, the same normalization approach was adapted to the blot level.

For the co-incubated CNR1/CNRIP1 antibody set, the membrane-level total-protein signal for each blot was divided by the highest membrane-level total-protein signal in the set: blot normalization factor = total-protein signal for blot X ÷ highest total-protein signal among blots in the antibody set. Each within-membrane normalized target signal was then divided by the corresponding blot normalization factor: final normalized signal = within-membrane normalized signal ÷ blot normalization factor. The Chameleon Duo ladder was used to identify molecular-weight positions and was not used as the normalization signal. Uncropped images of all immunoblots and corresponding total-protein stains used in these analyses are provided in Figure S1.

#### Statistical analyses and scientific rigor

Experimental subjects were assigned numeric identifiers and tail-color codes that concealed prenatal exposure history. Operant chamber assignments were randomized, and experimenters remained blinded to exposure history throughout behavioral testing, tissue collection, Western blotting, and protein quantification. Exposure assignments were verified after data collection by cross-referencing coded identifiers with the electronic colony record. Home-cage ethanol drinking measurements and the pre- and post-session syringe readings used to calculate ethanol intake during operant testing were recorded by hand. These records were subsequently transcribed independently by two experimenters, and the resulting datasets were electronically compared in Microsoft Excel to identify and resolve transcription discrepancies.

Statistical analyses were conducted using IBM SPSS Statistics version 29.0.2.0 and GraphPad Prism version 11. Prenatal alcohol exposure (ALC; present or absent) and prenatal cannabinoid exposure (CB; present or absent) were treated as separate factors in factorial analyses. Initial models included Sex, with sex-stratified analyses conducted to examine significant or trend-level effects involving Sex and to characterize previously established sex differences in ethanol-directed behavior. When necessary to identify differences among the four prenatal exposure groups, CON, ALC, CB, and ALC+CB were analyzed as levels of a single exposure factor. Significant or trend-level omnibus effects were followed by Šídák-adjusted comparisons unless otherwise specified. Potential outliers identified during initial assessment in SPSS were evaluated separately for each outcome using the ROUT procedure in GraphPad Prism with Q = 1%. Statistical significance was defined as p < .05, and .05 ≤ p < .10 was reported as a statistical trend. Pairwise comparisons, one-sample t tests, and correlation analyses were two-tailed.

Cage was the statistical unit for home-cage drinking analyses. Running cumulative ethanol intake across the 15-day assessment was analyzed using restricted maximum likelihood linear mixed-effects models with a random intercept for cage and Day modeled as a repeated factor using a first-order autoregressive covariance structure. The initial model included Sex, ALC, CB, Day, and their interactions. Follow-up models were conducted separately by sex using the factorial ALC × CB design and the four-level exposure factor. Ethanol preference was calculated for each cage as the natural logarithm of the ratio of cumulative ethanol-solution consumption to cumulative water consumption across the 10-day preference period. Preference was analyzed using a three-factor model containing Sex, ALC, CB, and their interactions, followed by sex-stratified factorial and four-level exposure analyses. Mean log-transformed preference was also compared with 0, representing equal ethanol and water consumption, using one-sample t tests with Holm correction across the four exposure groups within each sex.

Days required to meet operant-training criteria were analyzed using a three-factor model containing Sex, ALC, CB, and their interactions. Subjects that did not acquire the response criteria within the training period were assigned the maximum value of 15 days. Fixed-ratio 1 outcomes were analyzed using linear mixed-effects models with a random intercept for subject and Day modeled as a repeated factor. Models included Sex, ALC, CB, and their interactions, with sex-stratified and four-level exposure analyses conducted as appropriate. Changes in cumulative intake across FR1 testing were assessed using models that additionally included Day and its interactions. Associations among FR1 ethanol intake, active- and inactive-lever responding, and magazine entries were evaluated using Pearson correlations. Repeated-session mixed-effects regression models were also used to determine the simultaneous contributions of active-lever presses, inactive-lever presses, and magazine entries to ethanol intake. Variables were standardized before regression to obtain standardized coefficients.

Progressive-ratio ethanol intake, total ethanol-solution volume, and breakpoint were analyzed using mixed-effects models containing Sex, ALC, CB, ethanol concentration, and their interactions, with concentration treated as a repeated within-subject factor. Extinction responding across the three sessions was analyzed using mixed-effects models containing Sex, ALC, CB, Day, and their interactions. Normalized extinction responding was calculated as the mean number of active-lever presses during extinction divided by mean FR1 active-lever responding and multiplied by 100. Reinstatement ethanol intake and reinstatement intake normalized to mean FR1 intake were analyzed using three-factor models containing Sex, ALC, CB, and their interactions. Normalized extinction and reinstatement values were additionally compared with 100%, representing responding or intake equivalent to FR1 baseline, using one-sample t tests. Dunnett’s T3 comparisons were used in place of Šídák comparisons when homogeneity of variance was not supported.

CNR1 and CNRIP1 protein-expression values were transformed as ln(expression + 1) before analysis. Expression was initially evaluated using models containing Sex, ALC, CB, Region, and their interactions, with Region treated as a within-subject factor. Significant higher-order effects were examined using sex- and region-stratified mixed-effects models and Šídák-adjusted comparisons. Protein-behavior relationships were assessed separately by sex using multivariate general linear models. Separate models evaluated whether mPFC CNR1, mPFC CNRIP1, DMS CNR1, or DMS CNRIP1 expression was associated with the combined behavioral profile comprising mean FR1 ethanol intake, ethanol intake during the 40% progressive-ratio session, normalized extinction responding, and normalized reinstatement intake. Benjamini-Hochberg correction was applied across the four protein models within each sex. Significant multivariate associations were followed by Pearson correlations between protein expression and each behavioral outcome, with Benjamini-Hochberg correction applied across the four correlations. Spearman correlations were conducted as sensitivity analyses. Partial eta squared (ηp²) is reported as an effect-size estimate where applicable.

**Figure S1. Uncropped CNR1 and CNRIP1 immunoblots and corresponding total protein stains.** Full CNR1/CNRIP1 immunoblots and corresponding REVERT total protein stains are shown for each membrane. Lane annotations identify offspring sex and prenatal exposure group. Arrowheads indicate the bands quantified for CNR1 at approximately 53 kDa and CNRIP1 at approximately 20 kDa. Lanes indicated with * symbols were not included in analyses. CON, control; ALC, prenatal alcohol exposure; CB, prenatal cannabinoid exposure; ALC+CB, combined prenatal alcohol and cannabinoid exposure.

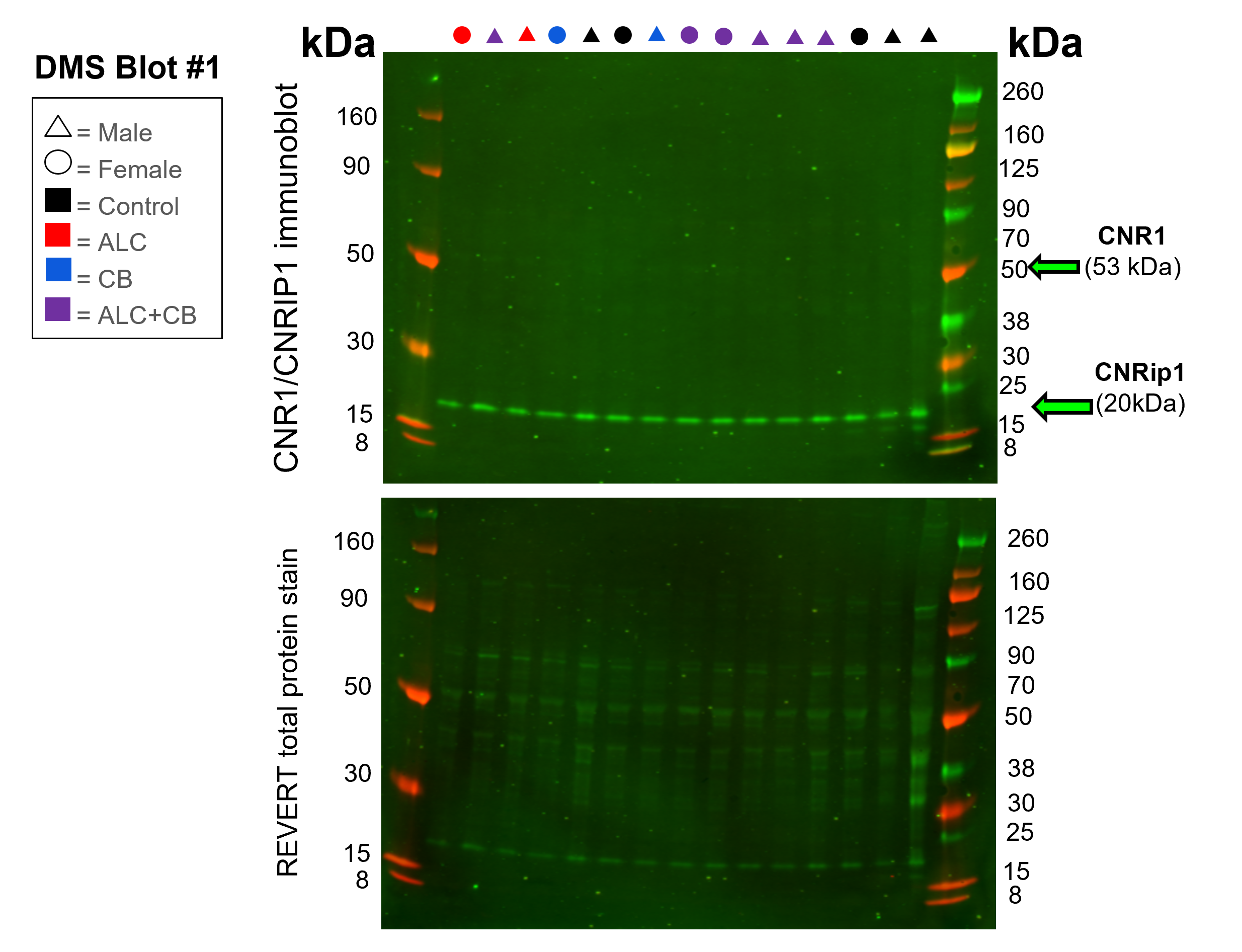

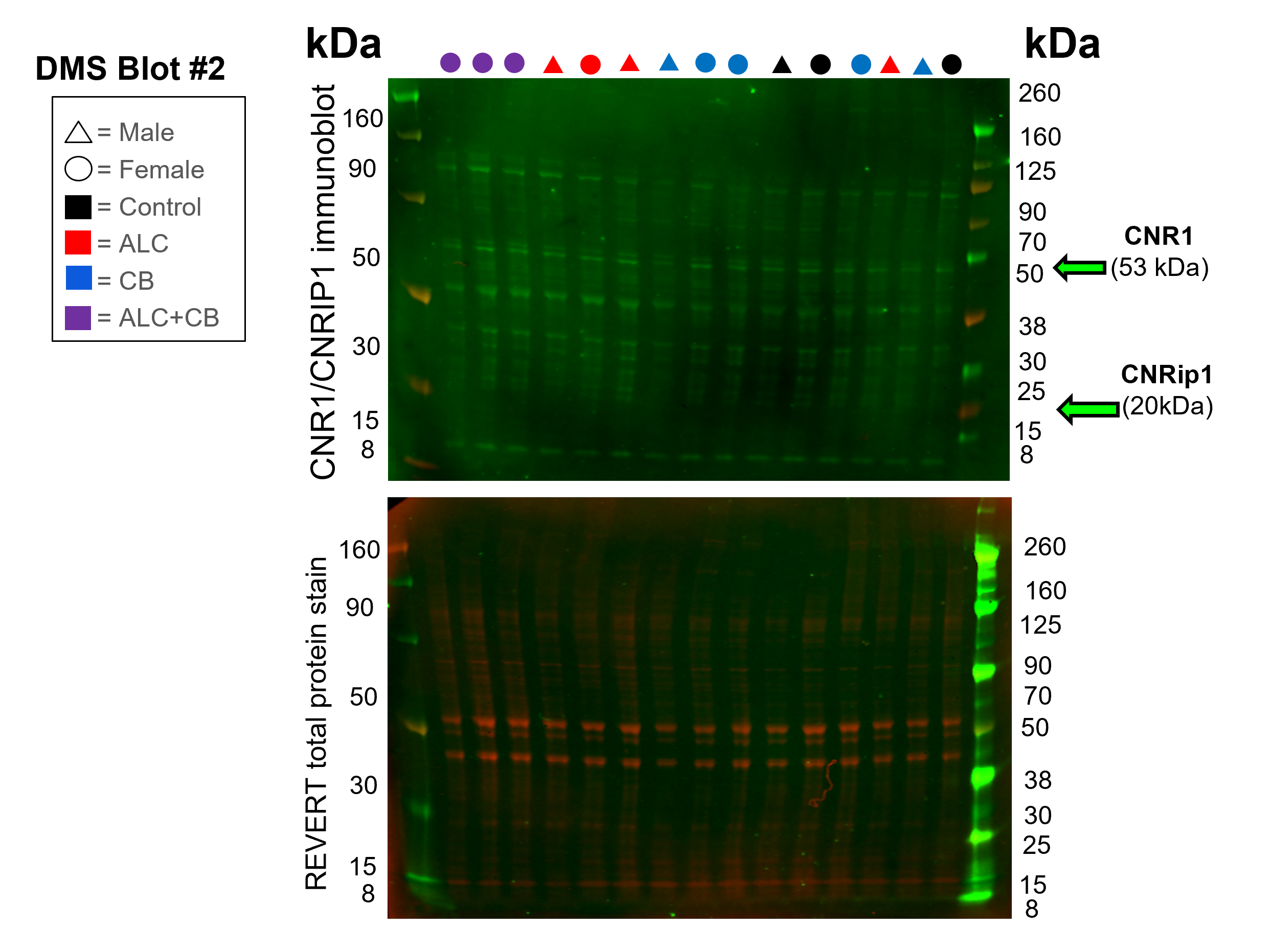

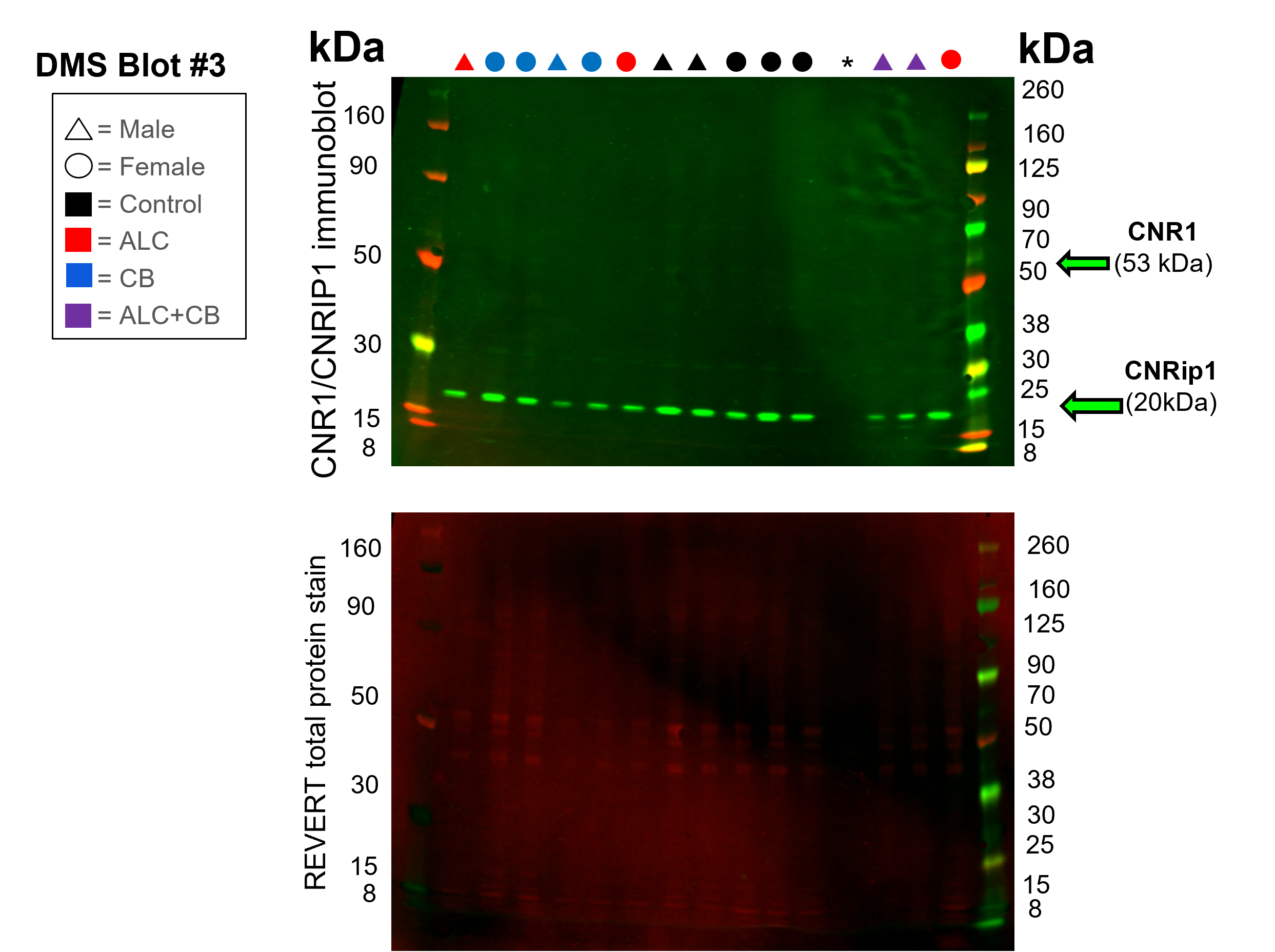

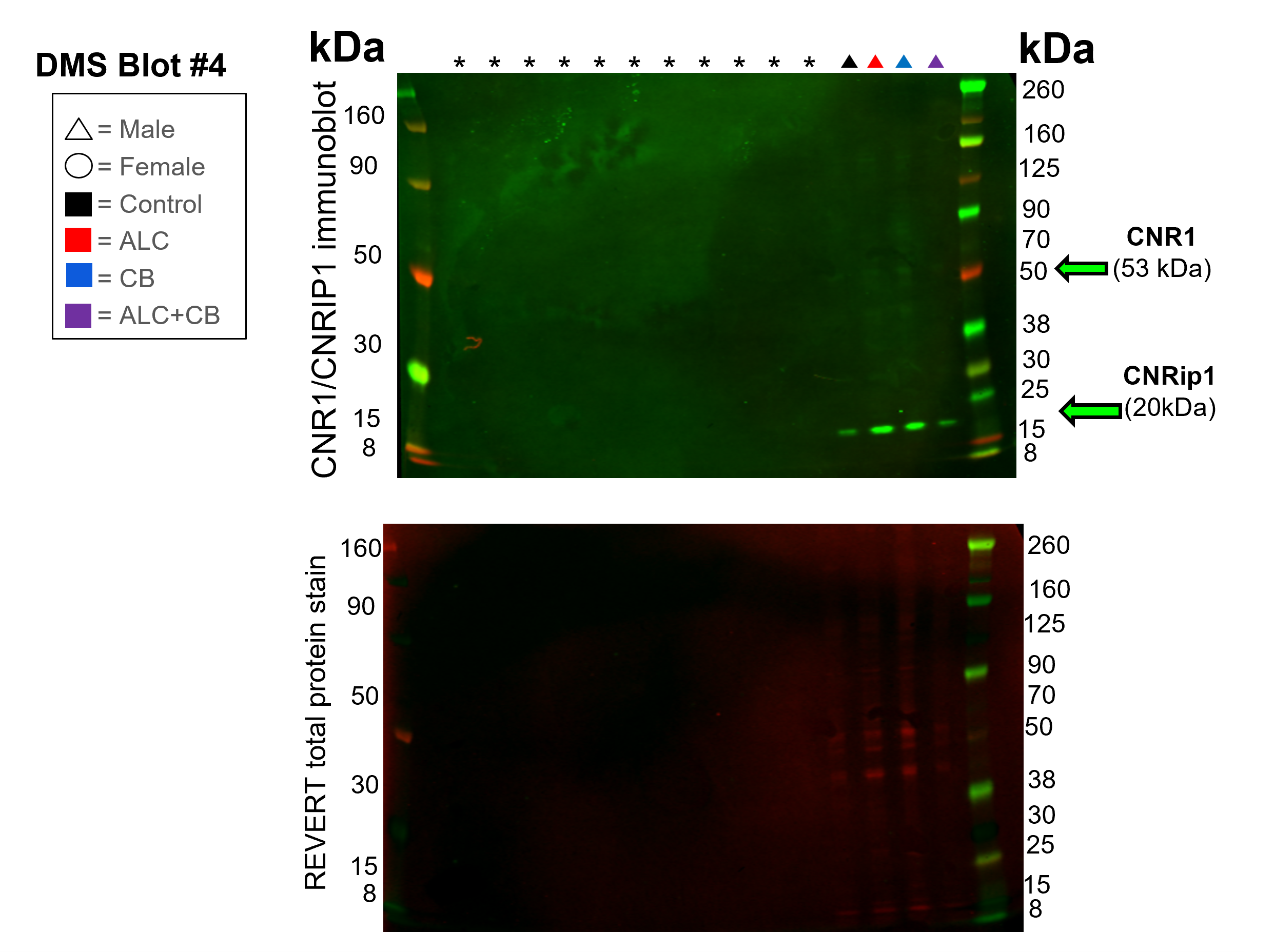

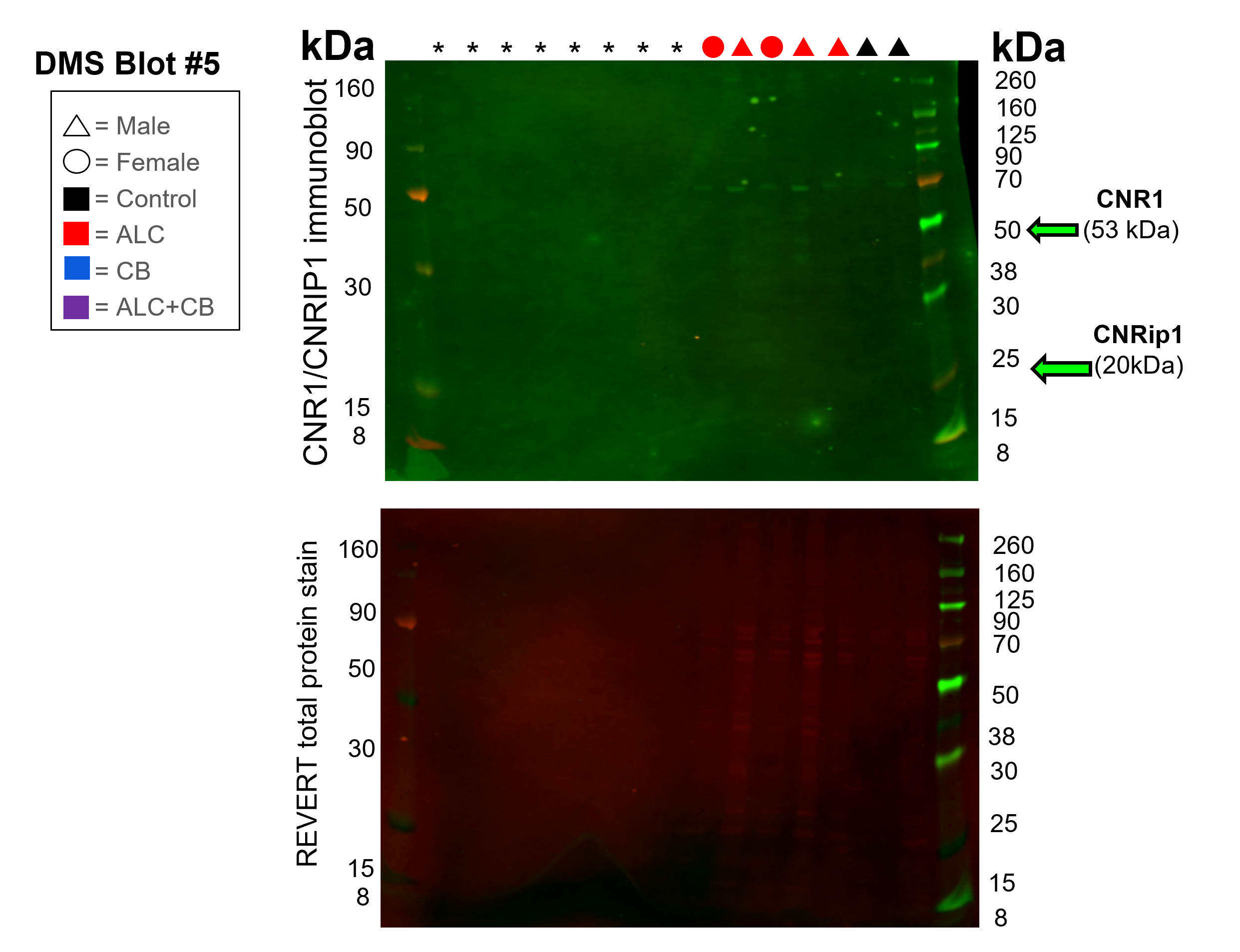

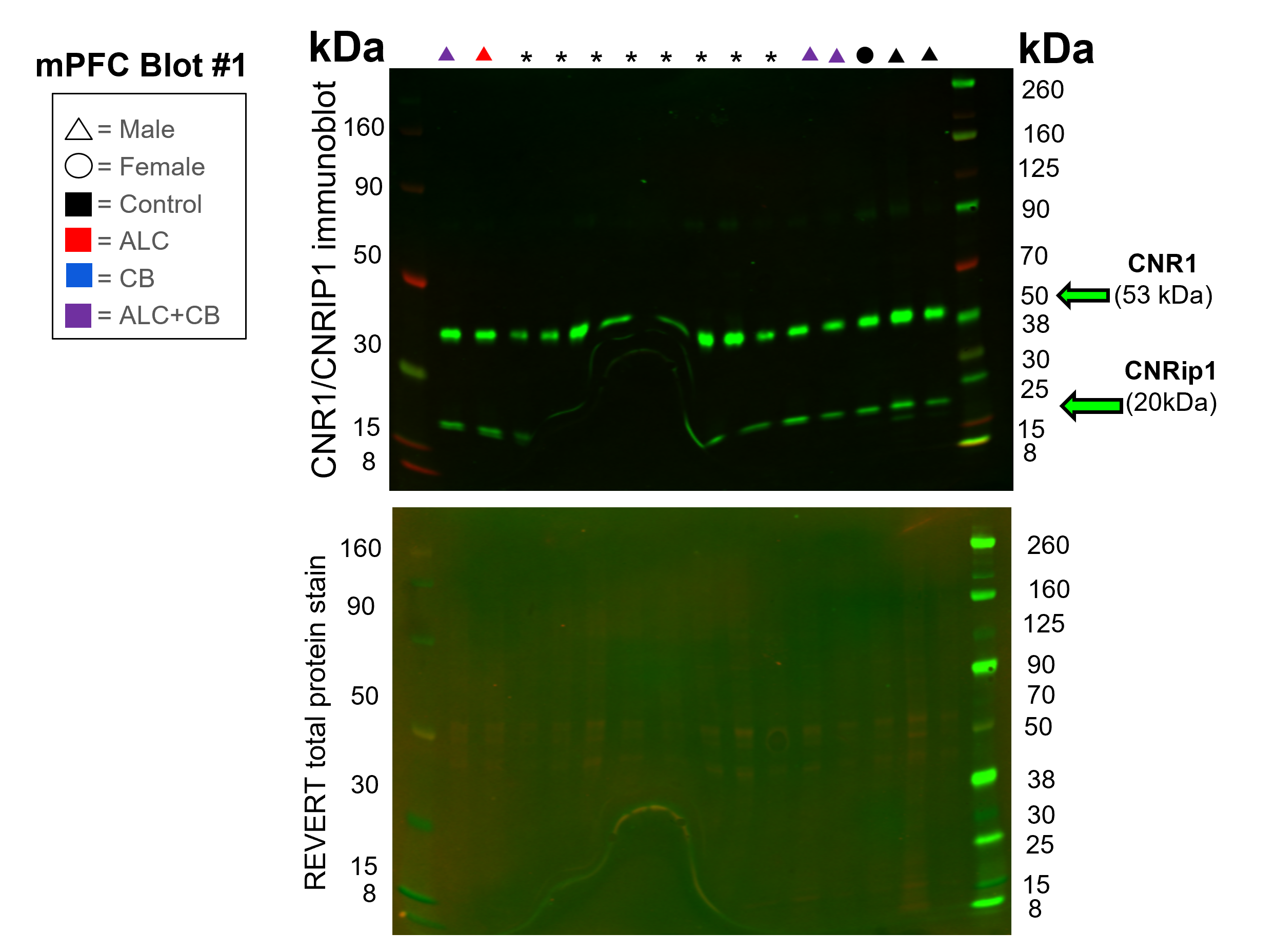

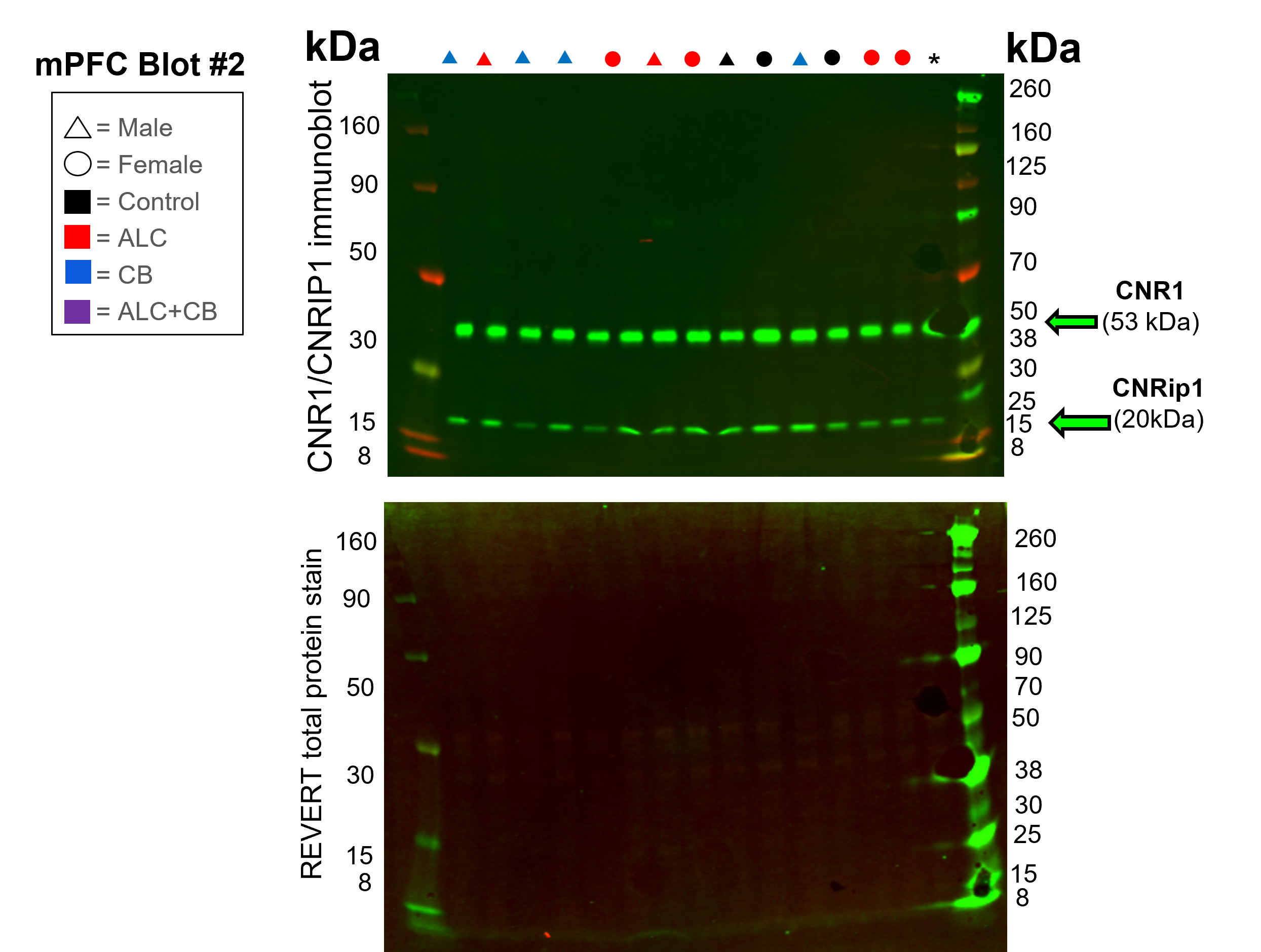

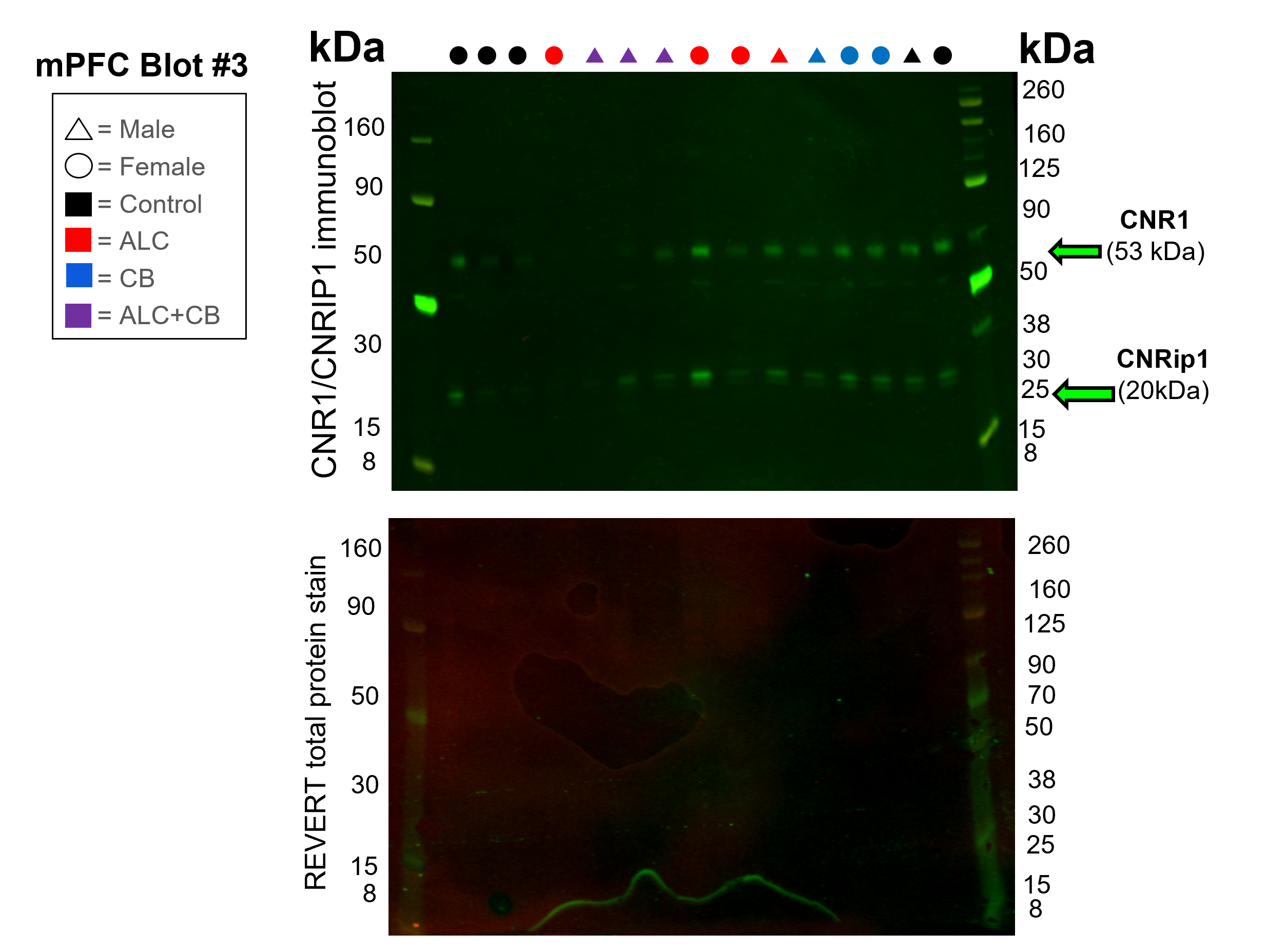

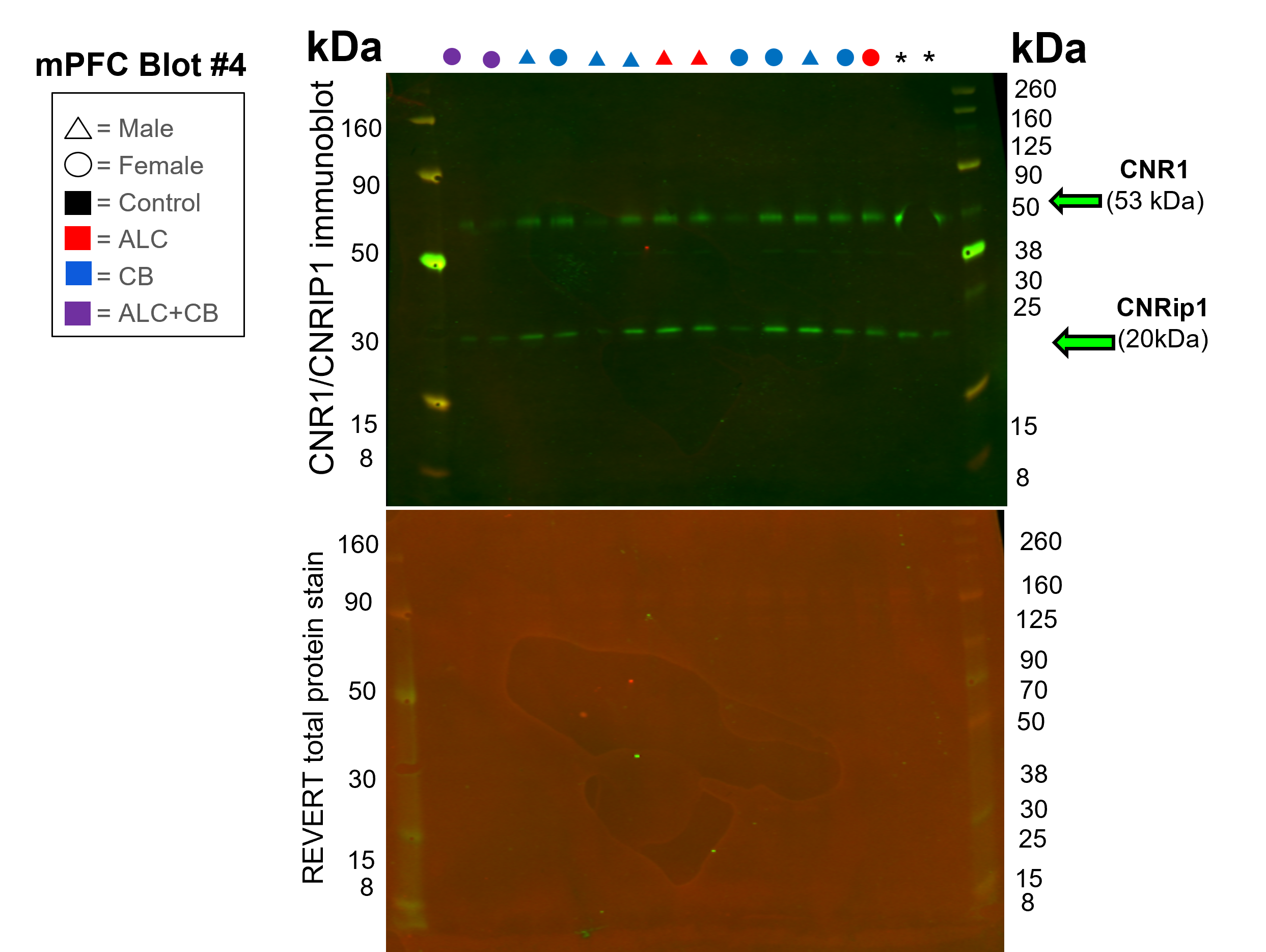

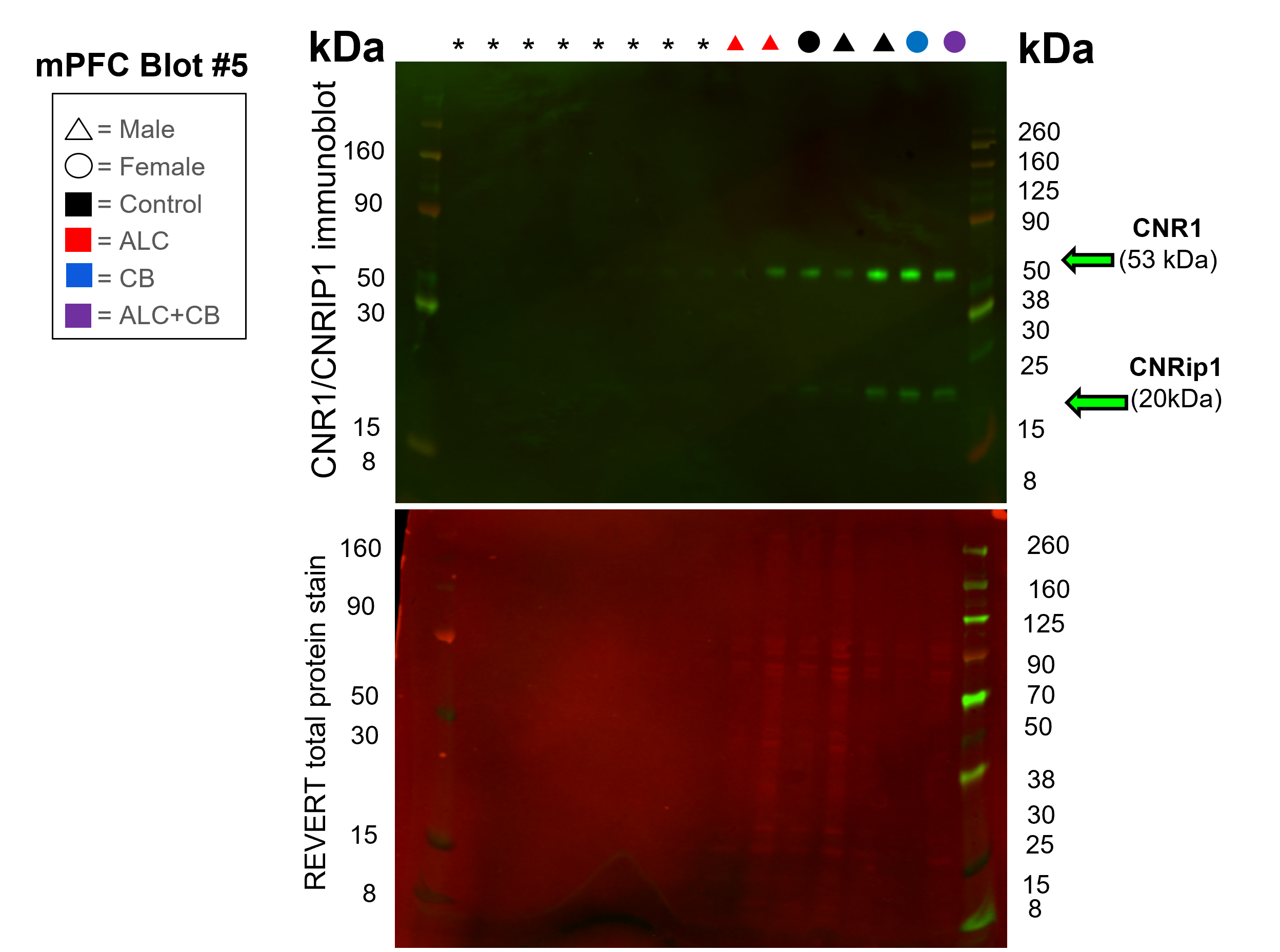

### Supplementary Results

*Prenatal cannabinoid exposure increases voluntary social drinking, particularly in male offspring.*

Home-cage drinking was assessed in male offspring from the CON (18 offspring, 8 cages), ALC (14 offspring, 6 cages), CB (14 offspring, 6 cages), and ALC+CB (8 offspring, 3 cages) groups and female offspring from the CON (18 offspring, 7 cages), ALC (15 offspring, 7 cages), CB (16 offspring, 7 cages), and ALC+CB (5 offspring, 2 cages) groups. During a 15-day, two-bottle choice drinking assessment in their home-cages, cumulative cage-level ethanol intake differed by sex, with female offspring consuming more ethanol than males independent of exposure, *[F*(1, 38.05) = 7.90, *p* = .008] (Fig. 1A). Any form of prenatal CB exposure (CB, ALC+CB = **CB+**; CON, ALC = **CB-**) produced a trend toward greater cumulative ethanol intake, *[F*(1, 38.05) = 3.42, *p* = .072] (Fig. 1B), and significantly interacted with Drinking Day, *[F*(14, 516.10) = 3.05, *p* < .001] (Table S1). Consistent with prior research in C57BL/6J mice [5, 6], control female offspring consumed significantly more ethanol per day than control male offspring, averaging 1.99 versus 1.10 g/kg/day, nested *t*(28) = 6.65, *p* < .001 (Fig. 1A). Given the overall sex difference in ethanol intake and previously established sex differences in drinking behavior, subsequent analyses were conducted separately in males and females.

Sex-stratified analyses indicated that prenatal exposure affected cumulative ethanol intake in males. CB+ males exhibited greater cumulative intake than CB- males, *[F*(1, 19.78) = 9.78, *p* = .005, ηp² = .331], and CB exposure interacted with Day, *[F*(14, 249.55) = 3.18, *p* < .001, ηp² = .151]. Prenatal ALC exposure also interacted with Day, *[F*(14, 249.55) = 2.26, *p* = .006, ηp² = .113], although comparisons between ALC-exposed (ALC, ALC+CB = **ALC+**) and non-ALC-exposed (CON, CB = **ALC-**) males were nonsignificant at every day (all Sidak-adjusted *ps* ≥ .091). All remaining exposure effects and interactions were nonsignificant (all *ps* ≥ .284; Table S2). Time-dependent comparisons indicated that CB+ males had greater cumulative intake than CB- males from Days 4 through 15 (all Sidak-adjusted *ps* ≤ .020). Comparisons among individual exposure groups further showed that CB-only males had greater cumulative intake than CON males from Days 7 through 15 (all *ps* ≤ .049; Fig. 1C). No comparisons involving ALC+CB males were significant (all *ps* ≥ .151).

In females, no exposure main effects or interactions reached significance. There was a trend toward a CB × Day interaction, *[F*(14, 265.09) = 1.60, *p* = .080, ηp² = .078], while all remaining exposure effects and interactions were nonsignificant (all *ps* ≥ .250; Table S3). In time-dependent comparisons, CB+ females had greater cumulative intake than CB− females on Day 15, mean difference = 9.39 g/kg, 95% CI [0.22, 18.56], *p* = .045. Although no significant exposure-group differences emerged from post hoc assessments (all Sidak-adjusted *ps* ≥ .345), Day 15 cumulative means suggest that this effect was driven by the ALC+CB group, whose cumulative intake was ~54% greater than the mean of the other three groups (43.63 vs. 28.33 g/kg; Fig. 1D).

*Males with prenatal ALC+CB exposure demonstrate greater preference for ethanol over water during two-bottle choice assessments.*

Ethanol preference was assessed in each exposure cage as the natural log of the daily ethanol-to-water consumption ratio, with 0 representing equal consumption of ethanol and water. Across the 10-day assessment, females exhibited higher ethanol-to-water ratios than males, *[F*(1, 38.19) = 9.89, *p* = .003, ηp² = .206], Fig. 1E–F; Table S4A). There was also a trend for prenatal CB exposure to increase ethanol-to-water ratios relative to no CB exposure, *[F*(1, 38.19) = 2.95, *p* = .094, ηp² = .072]. The ALC main effect and all interactions were nonsignificant (*ps* ≥ .111; Table S4A).

When analyzed separately by sex, males exhibited a trend toward higher ethanol-to-water ratios following prenatal CB exposure, *[F*(1, 19.04) = 4.26, *p* = .053, ηp² = .183]. No additional main effects, interactions, overall exposure-group effects, or Sidak-adjusted post hoc pairwise comparisons were significant (*ps* ≥ .111; Tables S4A–S5A). Finally, one-sample tests against equal liquid preference (0) indicated that ALC+CB males consumed proportionally more ethanol than water during two-bottle choice assessments, *t*(2) = 9.38, Holm-adjusted *p* = .044 (Fig. 1E; Table S5B). In contrast, CON males consumed proportionally more water [*t*(7) = −2.66, unadjusted *p* = .032] although this effect did not survive Holm correction (adjusted *p* = .096). No preference for either liquid was identified in the remaining male groups or in any female exposure group (Holm-adjusted *ps* ≥ .130; Fig. 1E–F; Table S5B).

*Prenatal alcohol and cannabinoid exposures do not alter overall acquisition of operant responding.*

To assess acquisition of operant responding for ethanol following social home-cage drinking, we examined the number of days required to reach the acquisition criterion (≥70% active-lever responding for three consecutive sessions). A three-way between-subjects ANOVA with Sex, ALC exposure, and CB exposure as factors revealed no main effects of Sex, ALC exposure, or CB exposure on days to criterion (all *ps* ≥ .769). However, prenatal ALC exposure differentially affected acquisition by sex, as indicated by a significant Sex × ALC interaction, *[F*(1, 88) = 6.74, *p* = .011, ηp² = .071]. Sidak-adjusted comparisons within each sex indicated that ALC+ males trended toward requiring more days to reach criterion than ALC- males (*p* = .059), whereas ALC+ females trended toward requiring fewer days than ALC- females (*p* = .080). Neither comparison reached statistical significance after adjustment. No interactions involving CB exposure were significant, including Sex × CB, ALC × CB, and Sex × ALC × CB (all *ps* ≥ .172; Table S6).

*Prenatal exposures increase operant ethanol consumption in female, but not male, offspring, with active lever responding predicting intake.*

Ethanol intake during the FR1 self-administration task was analyzed using a linear mixed-effects model with Sex, ALC exposure, and CB exposure as fixed factors and subject included as a random effect. Session was included as a repeated factor to account for within-subject variation across the three FR1 sessions (Fig. 2; Table S7). Averaged across sessions and exposure conditions, females consumed significantly more ethanol than males, *[F*(1, 85.54) = 7.04, *p* = .009, ηp² = .076]. Prenatal CB exposure tended to increase ethanol intake overall, *[F*(1, 85.54) = 2.96, *p* = .089, ηp² = .034], and this effect tended to differ by sex, Sex × CB: *[F*(1, 85.54) = 2.96, *p* = .089, ηp² = .033]. Sidak-adjusted simple-effects comparisons indicated that CB+ females consumed more ethanol than CB− females (*p* = .035), whereas CB+ and CB− males did not differ (*p* = .929; Fig. 2A). A significant Sex × ALC × CB interaction further indicated that the joint effects of prenatal ALC and CB exposure differed by sex, *[F*(1, 85.54) = 4.36, *p* = .040, ηp² = .049].

Sex-stratified analyses revealed no effects of prenatal exposure on FR1 ethanol intake in males (Fig. 2B). Neither the main effect of ALC, *[F*(1, 44.37) = 0.59, *p* = .446, ηp² = .013], nor CB, *[F*(1, 44.37) = 0.008, *p* = .930, ηp² < .001], was significant, and the ALC × CB interaction was also nonsignificant, *[F*(1, 44.37) = 1.78, *p* = .189, ηp² = .039]. In females, prenatal CB exposure increased ethanol intake during FR1 testing, *[F*(1, 41.45) = 4.09, *p* = .050, ηp² = .090]. Neither the ALC main effect, *[F*(1, 41.45) = 0.29, *p* = .592, ηp² = .007], nor the ALC × CB interaction, *[F*(1, 41.45) = 2.77, *p* = .103, ηp² = .063], was significant. Sidak-adjusted comparisons among the four exposure groups indicated that CB-only females consumed more ethanol than CON females (Fig. 2C; *p* = .015), while ALC-only females tended to consume more than CON females (*p* = .096). ALC+CB females did not differ from CON females (*p* = .228), and no comparisons among the three prenatally exposed groups were significant (all *ps* ≥ .817; Table S8).

To determine when exposure-group differences in cumulative intake emerged in females, cumulative ethanol intake was calculated through each of the three FR1 sessions, and Šídák-adjusted comparisons between each exposed group and CON were conducted within each day (Fig. 2C). No exposed group differed from CON on Day 1 (all *ps* ≥ .529). On Day 2, CB-only females tended to show greater cumulative ethanol intake than CON females (*p* = .071). By Day 3, ALC-only females also tended to show greater cumulative intake than CON females (*p* = .096), whereas CB-only females had significantly greater cumulative intake than CON females (*p* = .003). ALC+CB females did not differ from CON females on any day (all *ps* ≥ .312; Table S9).

In addition to ethanol intake, active and inactive lever presses and magazine entries were assessed during FR1 self-administration. The sex-combined linear mixed-effects model revealed a significant Sex × ALC × CB interaction for active lever presses, *[F*(1, 86.52) = 4.62, *p* = .034, ηp² = .051], indicating that the interaction between prenatal ALC and CB exposure differed by sex (Table S10A). Sex-stratified analyses revealed no main effects of ALC, *[F*(1, 45.98) = 0.86, *p* = .360, ηp² = .018], or CB, *[F*(1, 45.98) = 0.33, *p* = .570, ηp² = .007], in males, although the ALC × CB interaction showed a statistical trend, *[F*(1, 45.98) = 3.27, *p* = .077, ηp² = .066]. Descriptively, ALC+CB males exhibited the highest active-lever responding, whereas males exposed to ALC or CB alone exhibited lower mean responding than CON males (Table 1). In females, neither the main effect of ALC, *[F*(1, 39.68) = 0.51, *p* = .478, ηp² = .013], nor CB, *[F*(1, 39.68) = 1.45, *p* = .235, ηp² = .035], was significant, and there was no ALC × CB interaction, *[F*(1, 39.68) = 1.91, *p* = .175, ηp² = .046]. Nevertheless, all three exposed female groups exhibited descriptively greater mean active-lever responding than CON females (Table 1). These contrasting descriptive patterns are consistent with the significant Sex × ALC × CB interaction in the sex-combined model.

Inactive lever presses were analyzed to assess non-reinforced responding during FR1 self-administration. The linear mixed-effects model revealed a significant main effect of Sex, *[F*(1, 85.07) = 9.24, *p* = .003, ηp² = .098], with females making more inactive lever presses than males. There was no main effect of ALC exposure, *[F*(1, 85.07) = 0.05, *p* = .831, ηp² < .001]. The main effect of CB exposure showed a statistical trend, *[F*(1, 85.07) = 3.14, *p* = .080, ηp² = .036], with CB-exposed offspring making descriptively more inactive lever presses than offspring not exposed to CB. No interactions were significant, all *ps* ≥ .355 (Table S10B).

Magazine entries were analyzed as an index of engagement with the ethanol delivery port during FR1 self-administration. The linear mixed-effects model revealed a significant main effect of Sex, *[F*(1, 86.87) = 36.34, *p* < .001, ηp² = .295], with females making more magazine entries than males. There were no main effects of ALC exposure, *[F*(1, 86.87) = 1.42, *p* = .238, ηp² = .016], or CB exposure, *[F*(1, 86.87) = 0.02, *p* = .878, ηp² < .001], and no significant interactions, all *ps* ≥ .274 (Table S10C).

To assess whether subjects engaged in goal-directed alcohol-seeking behaviors during FR1 testing, lever discrimination was assessed using a discrimination index calculated as (active − inactive) / (active + inactive). Across all animals, discrimination was significantly greater than zero (*[F*(1, 87.99) = 19.20, *p* < .001]), indicating overall preference for the ethanol-associated lever among assessed offspring (Fig. 2D). Discrimination indices did not differ by Sex, ALC exposure, or CB exposure (all *ps* ≥ .11; Table S11), suggesting that prenatal exposures and offspring sex did not alter lever preference during operant ethanol self-administration.

To determine which operant behaviors were most closely associated with ethanol consumption, mean FR1 intake and operant-response measures were calculated across FR1 sessions for each subject. Ethanol intake was positively correlated with active-lever presses (Fig. 2E; Pearson’s *r* = .596, *p* < .001) and inactive-lever presses (Fig. 2F; Pearson’s *r* = .503, *p* < .001), but was not with magazine entries (Fig. 2G; Pearson’s *r* = .173, *p* = .095). Active-lever presses remained positively associated with ethanol intake after controlling for inactive-lever presses and magazine entries, partial *r* = .503, *p* < .001. Smaller independent associations were observed for inactive-lever presses, partial *r* = .238, *p* = .023, and magazine entries, partial *r* = .212, *p* = .043 (Table S12).

Consistent with the subject-level correlations, a mixed-effects regression accounting for repeated FR1 sessions within subjects identified active-lever presses as the strongest unique correlate of ethanol intake, standardized *β* = .688, 95% CI [.606, .770], *[F*(1, 222.68) = 273.82, *p* < .001]. Smaller positive associations were observed for magazine entries, standardized *β* = .115, 95% CI [.039, .192], *[F*(1, 212.27) = 8.77, *p* = .003], and inactive-lever presses, standardized *β* = .090, 95% CI [.005, .176], *[F*(1, 202.57) = 4.35, *p* = .038]. FR1 Day was not significantly associated with intake after accounting for these response measures, *[F*(2, 100.68) = 0.12, *p* = .887].

Together, these findings indicate that prenatal ALC or CB exposure increased operant ethanol consumption exclusively in female offspring. Across offspring, active-lever responding was more closely associated with ethanol intake than inactive-lever responding or magazine entries, supporting its use as the principal measure of ethanol-reinforced responding.

*Prenatal ALC and CB exposures sex-dependently increase motivation to obtain ethanol under a progressive ratio schedule.*

Next, ethanol intake during a progressive ratio task was analyzed using a linear mixed-effects model with Sex, ALC exposure, and CB exposure as fixed factors and subject included as a random effect, with ethanol concentration (20%, 30%, and 40%) modeled as a repeated factor to account for within-subject variation across sessions (Fig. 3; Table S13). Overall, ethanol intake increased as ethanol concentration increased (*[F*(2, 128.25) = 105.41, *p* < .001]), and females consumed more ethanol overall than males (*[F*(1, 88.17) = 12.43, *p* < .001]), with this sex difference varying across concentrations (*[F*(2, 128.25) = 5.45, *p* = .005]). Main effects of ALC (*[F*(1, 88.17) = 4.24, *p* = .043]) and CB (*[F*(1, 88.17) = 6.62, *p* = .012]) exposure were also observed. Both ALC (*[F*(2, 128.25) = 7.67, *p* < .001]) and CB (*[F*(2, 128.25) = 8.32, *p* < .001]) exposure interacted with ethanol concentration, indicating concentration-dependent effects of prenatal exposure on ethanol intake. A significant Sex × ALC × CB interaction was also observed (*[F*(1, 88.17) = 7.03, *p* = .010]), indicating that exposure effects differed between males and females.

In male offspring (Fig. 3A), ethanol intake varied as a function of both exposure and ethanol concentration (Table S14A). A significant ALC × CB interaction (*[F*(1, 46.05) = 6.78, *p* = .012]) indicated that co-exposure altered ethanol intake relative to single-drug exposures. Both ALC (*[F*(2, 67.09) = 7.41, *p* = .001]) and CB (*[F*(2, 67.09) = 3.15, *p* = .049]) also interacted with ethanol concentration. No exposure-group differences were observed at 20% or 30% ethanol (all *ps* ≥ .184; Table S14B). However, at 40%, ALC+CB male offspring consumed significantly more ethanol than control, ALC-only, and CB-only offspring (all *ps* < .001).

In female offspring (Fig. 3B), a significant CB × ethanol concentration interaction was observed (*[F*(2, 60.80) = 5.11, *p* = .009]; Table S15A), with no other exposure-related main effects or interactions (all *ps* ≥ .100). No exposure-group differences were observed at 20% or 30% ethanol (all *ps* ≥ .061; Table S15B). However, at 40%, ALC-only (*p* = .017), CB-only (*p* < .001), and ALC+CB (*p* = .016) offspring consumed more ethanol than controls, with no differences among the exposed groups (all *ps* ≥ .955).

To determine whether these effects reflected changes in total fluid consumption, the volume of liquid consumed during the progressive-ratio task was also analyzed (Table S16). Liquid consumption varied as a function of ethanol concentration (*[F*(2, 137.93) = 3.79, *p* = .025]), and significant interactions between ethanol concentration and both ALC exposure (*[F*(2, 137.93) = 4.06, *p* = .019]) and CB exposure (*[F*(2, 137.93) = 5.77, *p* = .004]) were observed. A significant Sex × ALC × CB interaction was also detected (*[F*(1, 87.97) = 5.23, *p* = .025]), indicating that the interactive effects of ALC and CB differed between males and females.

In male offspring, a significant ethanol concentration × ALC interaction was observed (*[F*(2, 71.71) = 3.79, *p* = .027]; Table S17A), with no other significant main effects or interactions (all *ps* ≥ .070). No exposure-group differences were observed at 20% or 30% ethanol (all *ps* ≥ .278; Table S17B). At 40%, ALC+CB offspring consumed more liquid than CB-only offspring (*p* = .032).

In female offspring, a significant CB × ethanol concentration interaction was observed (*[F*(2, 63.96) = 4.67, *p* = .013]; Table S18A), with no other exposure-related main effects or interactions (all *ps* ≥ .168). No exposure-group differences were observed at 20% or 30% ethanol (all *ps* ≥ .115; Table S18B). At 40%, CB-only offspring consumed more liquid than controls (*p* = .040), with a trend toward greater consumption in ALC-only offspring relative to controls (*p* = .080).

Breakpoint responding was then analyzed as an index of motivation to obtain ethanol (Table S19). Breakpoint increased with ethanol concentration (*[F*(2, 123.47) = 5.56, *p* = .005]), and a main effect of ALC exposure was observed (*[F*(1, 88.12) = 4.17, *p* = .044]). A significant CB × ethanol concentration interaction indicated that the effect of CB exposure varied across concentrations (*[F*(2, 123.47) = 9.56, *p* < .001]). A significant Sex × ALC × CB interaction further indicated that the interactive effects of ALC and CB differed between males and females (*[F*(1, 88.12) = 8.07, *p* = .006]).

In male offspring, a significant ALC × CB interaction indicated that the effect of each exposure on breakpoint responding depended on the presence of both ALC and CB exposures (*[F*(1, 46.63) = 6.09, *p* = .017]; Table S20A). A significant ALC × ethanol concentration interaction was also observed (*[F*(2, 53.29) = 3.93, *p* = .026]), with no other exposure-related effects reaching significance (all *ps* ≥ .051). Sidak-adjusted pairwise comparisons identified no exposure-group differences at 20%, 30%, or 40% ethanol (all *ps* ≥ .200; Table S20B).

In female offspring, a significant CB × ethanol concentration interaction indicated that the effect of CB exposure on breakpoint responding varied across concentrations (*[F*(2, 71.95) = 6.68, *p* = .002]; Table S21A), with no other exposure-related main effects or interactions (all *ps* ≥ .112). No exposure-group differences were observed at 20% or 30% ethanol (all *ps* ≥ .148; Table S21B). At 40%, CB-only offspring exhibited higher breakpoint responding than controls (*p* = .047), whereas no other exposure-group differences were observed (all *ps* ≥ .147).

Together, these findings indicate that prenatal ALC and CB exposure increase ethanol-directed behavior under high-demand conditions in a sex-dependent manner. Across measures, exposure-related differences were minimal at lower ethanol concentrations but emerged consistently at the highest concentration (40%), indicating that prenatal exposures primarily influence motivation for high-value ethanol rewards. Notably, CB+ exposure was more strongly associated with increased responding in females, whereas combined ALC+CB exposure produced more pronounced effects in males, demonstrating distinct sex-specific patterns in effort-based ethanol seeking.

*Prenatal ALC+CB co-exposure imposed persistent ethanol seeking during extinction in male offspring.*

Active lever presses during extinction were analyzed using a linear mixed-effects model with Sex, ALC exposure, and CB exposure as fixed factors, subject as a random effect, and extinction day (Days 1–3) as a repeated factor (Fig. 4A–D; Table S22). A main effect of ALC exposure was observed (*[F*(1, 89.96) = 5.09, *p* = .027]), along with significant Sex × CB (*[F*(1, 89.96) = 4.01, *p* = .048]) and Sex × ALC × CB (*[F*(1, 89.96) = 10.25, *p* = .002]) interactions, indicating that exposure effects on extinction responding differed between males and females. No effects involving extinction day were detected (all *ps* ≥ .249), indicating that differences in responding reflect overall levels of ethanol-seeking persistence rather than altered rates of extinction.

In male offspring, a main effect of ALC exposure (*[F*(1, 46.80) = 4.92, *p* = .032]) and a significant ALC × CB interaction (*[F*(1, 46.80) = 6.04, *p* = .018]) were observed (Fig. 4A; Table S23A). The main effect of CB exposure showed a trend (*[F*(1, 46.80) = 3.53, *p* = .066]), while no effects involving extinction day were detected (all *ps* ≥ .105). Sidak-adjusted pairwise comparisons, collapsed across extinction days, revealed that ALC+CB offspring exhibited greater active lever responding than ALC-only (*p* = .033) and CB-only (*p* = .021) offspring (Table S23B). Neither single-exposure group differed from controls (all *ps* ≥ .996), while ALC+CB offspring showed a trend toward greater responding than controls (*p* = .066).

In female offspring, a significant ALC × CB interaction was observed (*[F*(1, 41.80) = 7.33, *p* = .010]; Fig. 4C; Table S24A), indicating that the effect of each exposure on extinction responding depended on the presence of the other. No main effects of ALC or CB exposure were detected (all *ps* ≥ .392), and no effects involving extinction day were observed (all *ps* ≥ .160). Although ALC-only offspring showed a trend toward greater active lever responding than controls (*p* = .079), no Sidak-adjusted pairwise comparison reached significance (all *ps* ≥ .079; Table S24B).

To account for individual differences in reinforced responding, extinction active lever pressing was expressed relative to each subject’s FR1 responding (Fig. 4B, D; Table S25). Analysis of extinction responding relative to baseline revealed a significant main effect of ALC exposure (*[F*(1, 85.89) = 5.20, *p* = .025]), as well as significant Sex × ALC (*[F*(1, 85.89) = 6.90, *p* = .010]), Sex × CB (*[F*(1, 85.89) = 5.65, *p* = .020]), and Sex × ALC × CB (*[F*(1, 85.89) = 9.15, *p* = .003]) interactions. The three-way interaction indicated that the interactive effects of ALC and CB on normalized extinction responding differed between males and females.

In male offspring, a significant main effect of ALC exposure (*[F*(1, 44.08) = 12.98, *p* < .001]) and an ALC × CB interaction (*[F*(1, 44.08) = 6.86, *p* = .012]) were observed, while the main effect of CB exposure showed a trend (*[F*(1, 44.08) = 3.27, *p* = .077]; Table S26A). Šídák-adjusted comparisons showed that ALC+CB offspring exhibited greater extinction responding relative to baseline than control (*p* = .0059), ALC-only (*p* = .0229), and CB-only (*p* = .0023) offspring (Fig. 4B; Table S26B). No differences were observed among the control, ALC-only, and CB-only groups (all *ps* ≥ .288).

In female offspring, the ALC × CB interaction showed a trend (*[F*(1, 40.80) = 3.43, *p* = .071]), while neither main effect approached significance (all *ps* ≥ .121; Table S27A). No Šídák-adjusted exposure-group comparison reached significance (all *ps* ≥ .187; Fig. 4D; Table S27B).

To evaluate whether responding was suppressed relative to prior reinforced levels, one-sample *t*-tests compared extinction responding relative to FR1 baseline with a theoretical value of 100%. In male offspring, control, ALC-only, and CB-only groups exhibited extinction responding significantly below baseline (all *ps* < .0001), whereas ALC+CB offspring did not differ from baseline (*p* = .597; Table S26B). Thus, ALC+CB was the only male exposure group that did not show evidence of reduced responding relative to its reinforced baseline. In females, all exposure groups exhibited extinction responding significantly below baseline (all *ps* ≤ .0005; Table S27B), indicating suppression of responding relative to reinforced levels regardless of prenatal exposure history.

Together, these findings indicate that prenatal ALC+CB exposure selectively increased extinction responding relative to reinforced baseline in male offspring. This effect remained evident after responding was scaled to each subject’s FR1 baseline and was not observed in females. Combined with the absence of exposure-related differences in responding across extinction days in the non-normalized analysis, these results support persistently elevated ethanol seeking across extinction sessions in male ALC+CB offspring, without evidence of a different day-to-day extinction trajectory.

*Prenatal ALC+CB exposure increased reinstatement ethanol intake relative to baseline in male offspring.*

Ethanol intake during a one-day reinstatement assessment was analyzed using a three-way ANOVA with Sex, ALC exposure, and CB exposure as between-subject factors (Fig. 4E, G; Table S28A). This analysis revealed significant main effects of ALC exposure, *[F*(1, 87) = 4.26, *p* = .042], and CB exposure, *[F*(1, 87) = 4.11, *p* = .046], as well as a significant Sex × ALC × CB interaction, *[F*(1, 87) = 5.55, *p* = .021]. The three-way interaction indicated that prenatal exposure effects on reinstatement ethanol intake differed between male and female offspring. In male offspring, there was a main effect of CB exposure, *[F*(1, 45) = 5.07, *p* = .029], and an ALC × CB interaction, *[F*(1, 45) = 5.23, *p* = .027] (Fig. 4E; Table S28B). However, Dunnett T3-adjusted pairwise comparisons did not identify significant differences between exposure groups (all *ps* ≥ .505). In female offspring, there were no main effects of ALC or CB exposure and no ALC × CB interaction (all *ps* ≥ .183; Fig. 4G; Table S28C).

To account for within-subject differences in reinforced ethanol intake, we expressed reinstatement ethanol intake relative to FR1 baseline (Fig. 4F, H; Table S29A). This analysis revealed a significant Sex × CB interaction, *[F*(1, 87) = 4.90, *p* = .030], indicating that the effect of prenatal CB exposure differed between male and female offspring. In males, we observed a main effect of CB exposure, *[F*(1, 45) = 6.44, *p* = .015], as well as trend-level effects of ALC exposure, *[F*(1, 45) = 3.33, *p* = .075], and the ALC × CB interaction, *[F*(1, 45) = 3.11, *p* = .084] (Fig. 4F; Table S29B). Šídák-adjusted comparisons showed that ALC+CB offspring exhibited greater reinstatement intake relative to baseline than CON and ALC offspring (both *ps* = .039). No other comparisons were significant (all *ps* ≥ .124). In females, we detected no main effects of ALC or CB exposure and no ALC × CB interaction (all *ps* ≥ .538; Fig. 4H; Table S29C).

To determine whether reinstatement intake differed from prior reinforced drinking, we performed two-tailed one-sample *t* tests comparing reinstatement ethanol intake relative to FR1 baseline against 100%. In males, CON, ALC, and CB offspring exhibited reinstatement intake significantly below baseline (all *ts* ≥ 2.98, all *ps* ≤ .011), whereas ALC+CB offspring did not differ from baseline, *t*(7) = 0.25, *p* = .808 (Table S29B). In females, CON, ALC, and CB offspring also exhibited reinstatement intake significantly below baseline (all *ts* ≥ 3.67, all *ps* ≤ .003). ALC+CB offspring showed a trend toward below-baseline intake, *t*(4) = 2.40, *p* = .075, with mean intake remaining at 61.71% of baseline and comparable to the other female exposure groups (51.00–58.63%; Table S29C).

*Prenatal ALC and CB exposures produce sex- and brain region-specific changes in CNR1 and CNRIP1 expression.*

CNR1 and CNRIP1 protein expression (Fig. 5A) was analyzed using a four-way ANOVA with Sex, ALC exposure, CB exposure, and Region as fixed factors (Table S30). CNR1 levels were significantly higher in the DMS than the mPFC overall, *[F*(1, 47.76) = 20.54, *p* < .001]. The effect of CB exposure varied by region, *[F*(1, 47.76) = 7.65, *p* = .008], as did the combined effects of ALC and CB exposure, *[F*(1, 47.76) = 9.09, *p* = .004]. The region-dependent effect of CB exposure further differed by sex, *[F*(1, 47.76) = 5.84, *p* = .019]. The ALC × Region × Sex interaction, *[F*(1, 47.76) = 3.22, *p* = .079], and the ALC × CB × Region × Sex interaction, *[F*(1, 47.76) = 3.13, *p* = .083], reached trend-level significance. Within the same samples, CNRIP1 protein expression was analyzed using the same four-way model (Table S31). CNRIP1 levels were significantly higher in the DMS than the mPFC overall, *[F*(1, 47.48) = 174.82, *p* < .001]. ALC and CB exposure interacted to alter CNRIP1 expression across sexes and regions, *[F*(1, 47.55) = 6.34, *p* = .015]. The ALC × Sex interaction also reached trend-level significance, *[F*(1, 47.55) = 3.89, *p* = .055].

To characterize protein-expression patterns within each sex and region, CNR1 and CNRIP1 were analyzed jointly using mixed-effects models with Protein as a matched factor and ALC and CB exposure as fixed factors (Fig. 5B). In the male mPFC, there were significant main effects of Protein, *[F*(1, 49) = 9.87, *p* = .003], and ALC exposure, *[F*(1, 49) = 4.24, *p* = .045], as well as a trend-level Protein × ALC interaction, *[F*(1, 49) = 3.73, *p* = .059] (Fig. 5C; Table S32A). Šídák-adjusted comparisons showed that ALC+CB offspring exhibited higher CNR1 expression than CON, ALC, and CB offspring (all *ps* ≤ .010). No exposure-group differences were detected for CNRIP1 (all *ps* > .9999; Table S32B).

In the male DMS, there were a significant main effect of Protein, *[F*(1, 19) = 4.71, *p* = .043], and an ALC × CB interaction, *[F*(1, 27) = 6.43, *p* = .017] (Fig. 5D; Table S33A). The Protein × CB interaction, *[F*(1, 19) = 3.82, *p* = .066], and Protein × ALC × CB interaction, *[F*(1, 19) = 3.09, *p* = .095], reached trend-level significance. Šídák-adjusted comparisons showed that ALC offspring exhibited higher CNRIP1 expression than CON and ALC+CB offspring (all *ps* ≤ .016). No exposure-group differences were detected for CNR1 (all *ps* ≥ .977; Table S33B).

In the female mPFC, there were significant main effects of Protein, *[F*(1, 23) = 6.39, *p* = .019], and CB exposure, *[F*(1, 24) = 4.41, *p* = .046], as well as a trend-level Protein × CB interaction, *[F*(1, 23) = 3.98, *p* = .058] (Fig. 5E; Table S34A). Šídák-adjusted comparisons showed that CB and ALC+CB offspring exhibited lower CNR1 expression than CON offspring (both *ps* ≤ .045). No other exposure-group differences were detected for CNR1 (all *ps* ≥ .300), and CNRIP1 did not differ among exposure groups (all *ps* > .9999; Table S34B).

Finally, in the female DMS, there were no significant main effects or interactions on protein expression (all *ps* ≥ .106; Fig. 5F; Table S35).

*mPFC CNR1 expression was associated with ethanol-directed behaviors in male offspring.*

Given previously identified sex differences in ethanol-directed behaviors and protein expression, protein-behavior associations were evaluated separately in male and female offspring. Sex-stratified multivariate general linear models assessed whether each region-specific protein measure was associated with a combined behavioral profile comprising average FR1 ethanol intake, 40% progressive-ratio ethanol intake, normalized extinction responding, and normalized reinstatement responding (Fig. 6A; Table S36). Among the four region-specific protein measures examined, mPFC CNR1 was the only measure associated with the combined behavioral profile. This association was significant in males (Pillai’s trace = .353, *[F*(4, 22) = 3.01, *p* = .040, partial η² = .353]) but not females (*p* = .392). However, the male omnibus association did not survive Benjamini-Hochberg correction across the four region-specific protein measures (*q* = .160).

In behavior-specific correlations, higher log-transformed mPFC CNR1 expression in males was positively associated with average FR1 ethanol intake (*r*(26) = .404, *p* = .033), 40% progressive-ratio ethanol intake (*r*(26) = .498, *p* = .007; Fig. 6B), normalized extinction responding (*r*(26) = .483, *p* = .009; Fig. 6B), and normalized reinstatement responding (*r*(25) = .501, *p* = .008; Table S37). All four Pearson correlations remained significant after Benjamini-Hochberg correction across the four behavioral outcomes (*q*s = .012-.033). Spearman sensitivity analyses yielded significant associations with progressive-ratio intake (ρ = .450, *p* = .016), normalized extinction responding (ρ = .461, *p* = .013), and normalized reinstatement responding (ρ = .421, *p* = .029), whereas the association with FR1 ethanol intake was weaker (ρ = .322, *p* = .095).

**Supplemental References**

1.	Rouzer, S.K., et al., *Early life outcomes of prenatal exposure to alcohol and synthetic cannabinoids in mice.* Drug and Alcohol Dependence Reports, 2025. **16**: p. 100356.

2.	Godynyuk, E., et al., *An Open-Source, Automated Home-Cage Sipper Device for Monitoring Liquid Ingestive Behavior in Rodents.* eneuro, 2019. **6**(5): p. ENEURO.0292-19.2019.

3.	Blegen, M.B., et al., *Alcohol operant self-administration: Investigating how alcohol-seeking behaviors predict drinking in mice using two operant approaches.* Alcohol, 2018. **67**: p. 23-36.

4.	Livy, D.J., S.E. Parnell, and J.R. West, *Blood ethanol concentration profiles: a comparison between rats and mice.* Alcohol, 2003. **29**(3): p. 165-71.

5.	Rivera-Irizarry, J.K., et al., *Sex differences in binge alcohol drinking and the behavioral consequences of protracted abstinence in C57BL/6J mice.* Biology of sex differences, 2023. **14**(1): p. 83.

6.	Sneddon, E.A., R.D. White, and A.K. Radke, *Sex differences in binge‐like and aversion‐resistant alcohol drinking in C57 BL/6J mice.* Alcoholism: clinical and experimental research, 2019. **43**(2): p. 243-249.
